# Global protein expression profiling in stem cell factor stimulated human Acute megakaryoblastic leukemia cells identifies CFL1, GSN and CCT8 as prognostic biomarkers for Acute Myeloid Leukemia

**DOI:** 10.64898/2026.08.24.746695

**Authors:** Anand Kuttanparambil Ravi, Gopika Gopan, Sudhakar Arumugam, Aiswarya Sethumadhavan, Maheswaran Mani

## Abstract

**Background:** The stem cell factor receptor or c-Kit is a type III receptor tyrosine kinase, activated by its ligand Stem cell factor (SCF). Up on activation, c-kit induces signaling pathways that regulates blood cell proliferation, survival, differentiation, and migration. Several studies reported that c-Kit/SCF signaling, contributes to the development and progression of acute myeloid leukemia (AML) in patients. However, the downstream proteins regulated by c-kit activation and their clinical significance in AML remain poorly explored.

**Methods:** Human Acute megakaryoblastic leukemia (Mo7e) cells, were-stimulated with SCF and global protein expression were profiled using two-dimensional gel electrophoresis coupled with MALDI-TOF and LC-MS/MS. Differentially expressed proteins were functionally characterized and validated using patient data from the TCGA-LAML and matched normal data from GTEx, GEO datasets, and quantitative RT-PCR. Their diagnostic and prognostic significance was assessed using ROC, Cox regression, LASSO, Kaplan Meier survival analyses, and a prognostic nomogram model.

**Results:** Proteomic profiling identified 14 differentially expressed proteins in SCF-stimulated Mo7e cells, which are predicted to involved in cytoskeletal organization, protein folding, metabolism, vesicular trafficking, and translational regulation. Transcriptomic analysis of the TCGA-LAML cohort revealed significant dysregulation of CFL1, CCT8, HSP90B1, MDH2, EIF5A, GSN, and TPI1. Integrated ROC, Cox regression, and LASSO analyses identified CFL1, CCT8, and GSN as the most robust prognostic biomarkers associated with poor overall survival in LAML patients. Their expression patterns were validated in independent GEO datasets and by qRT-PCR in SCF stimulated Mo7e cells. Finally, a three-gene nomogram model was developed and validated to predict the overall survival probability of AML patients at 1-, 3-, and 5-year time points.

**Conclusions:** This study identifies CFL1, CCT8, and GSN as key downstream effectors of c-Kit signaling as prognostic biomarkers for AML. These findings provide mechanistic insights into c-Kit-driven leukemogenesis and establish a clinically relevant three-gene signature for AML risk stratification and potential therapeutic targeting.

**Graphical Abstract:** 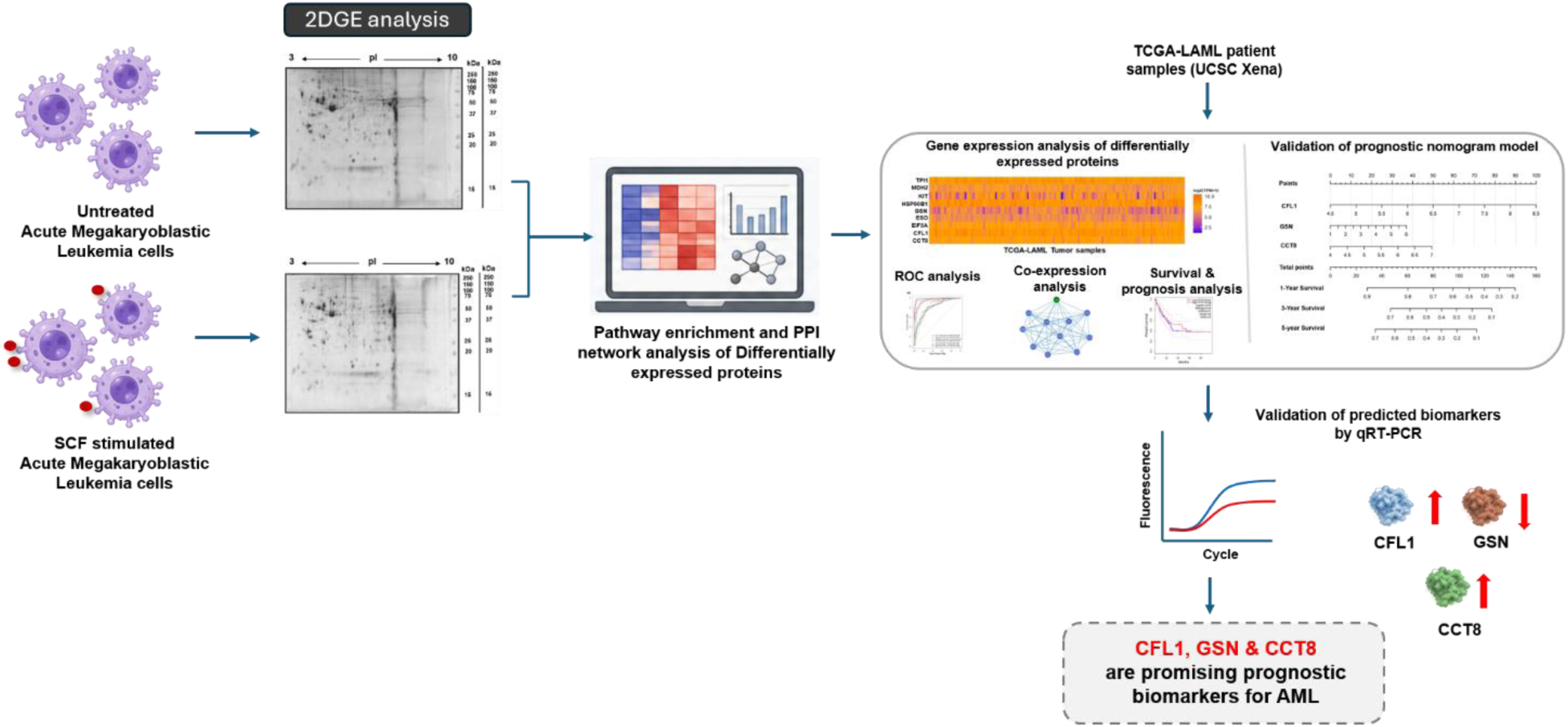

## Introduction

Stem cell factor (SCF), also known as Kit ligand (KITLG), mast cell growth factor or steel factor is a multifunctional cytokine encoded by the Steel (Sl) locus, which is essential for the regulation of hematopoiesis, melanogenesis and germ cell development. SCF is the physiological ligand for c-Kit (CD117), a type III receptor tyrosine kinase that is primarily expressed on blood forming stem and progenitor cells [1]. Following SCF binding, c-Kit homodimerizes and subsequently intracellular tyrosine residues undergo autophosphorylation. This triggers different downstream signaling pathways including mTOR, JAK-STAT, PI3K-AKT and MAPK-ERK cascades [2]. Together these pathways regulate key cellular functions such as cell proliferation, differentiation, survival, migration, apoptosis, and cell cycle progression [3]. Overall, the SCF/c-Kit signaling axis is critically involved in hematopoietic cells homeostasis.

c-Kit signaling has been involved in numerous cancers including acute myeloid leukemia. c-Kit is highly overexpressed in leukemic blasts and generally correlates with immature leukemic characteristics [4]. Moreover, KIT-activating mutations are frequently reported in some subtypes of acute myeloid leukemia (AML), especially core-binding factor AML (CBF-AML), where they correspond with higher risk of relapse and adverse clinical outcome [5]. Altogether, c-Kit signaling increases leukemic cell proliferation, decreased apoptosis, increased self-renewal potential and disease progression. These leukemic blasts suppress the normal hematopoiesis leading to anemia, thrombocytopenia, recurrent infections and bone marrow failure [6]. Therefore, the investigation of prognostic biomarkers in leukemogenesis is a clinical priority.

The c-Kit signaling activates numerous protein that have been involved in AML development, survival, stemness, and treatment resistance, including MYC [7], CCND1 [8], STAT5 [9], AKT1[10], CXCR4[11], HOXA9[12], MEIS1[13], and RUNX1[14]. Therefore, characterization of c-Kit-responsive proteins may offer substantial insights into the molecular mechanisms underlying AML progression and help to identify prognostic biomarkers and treatment options. Recently, high-throughput proteomics has evolved to the point of large-scale profiling of changes in protein expression in leukemic cells. However, there are few studies that discuss c-Kit activated proteomic profile and its prognostic importance in AML. In this study, we investigated the global protein changes in SCF-stimulated leukemia cells. We used human megakaryoblastic leukemia cell line (M07e) as an experimental model, as it has strong c-Kit expression and reproducible response to SCF stimulation [15]. Furthermore, the clinical importance of differentially expressed proteins was assessed using publicly available datasets of AML patients.

## Materials and methods

### Stimulation Mo7e cells

The human megakaryocytic cell line Mo7e (ACC 104) was obtained from the German Collection of Microorganisms and Cell Cultures (DSMZ, Germany). Cells were maintained in RPMI 1640 medium (Gibco, USA) supplemented with 10% fetal calf serum (HiMedia, India) and recombinant human IL-3 (20 ng/mL; Invitrogen, USA). Prior to treatment, the cells were deprived of growth factors for 12 h and subsequently stimulated with 100 ng/mL stem cell factor (SCF; PeproTech Asia, Israel).

### Protein expression profiling by 2-Dimensional Gel Electrophoresis

SCF-stimulated (100 ng/mL) and non-stimulated Mo7e cells were subjected to protein extraction and two-dimensional gel electrophoresis following the previously described methodology, with modifications where applicable [16]. Cells were lysed in 1 ml of urea lysis buffer containing 30 mM Tris pH 8.5 (Himedia, India), 2 M Thiourea, 7 M Urea, 4% CHAPS (w/v), 1% IPG buffer (pH: 3-10), 2% DTT, 1 mM sodium orthovanadate, 1 mM β-Glycerophosphate, and 1 mM PMSF (Sigma Aldrich, USA). As recommended by the manufacturer, protease inhibitor (Complete; Roche, Switzerland) was added. The Bradford assay was used to measure the amount of protein in the supernatant after cell extracts were centrifuged for one hour at 14000 rpm. After loading 450 μl of destreak rehydration solution (GE Healthcare, USA) containing 900 μg of protein onto immobiline dry strips (24 cm, pH 3-10), the strips were left to rehydrate for eighteen hours. The EttanIPGphor 3 IEF system (GE Healthcare, USA) was used to perform isoelectric focusing at 20°C. Isoelectric Focusing (57300 Vh) was performed at 500 V for two hours, followed by a gradient rise to 1000 V (1000 Vh), 8000 V (20000 Vh), and 10000 V (25000 Vh), 10000 V for one hour, and 100 V for the last hour.

IPG strips were equilibrated for the second-dimension separation following isoelectric focusing. In short, IPG strips were incubated for 15 minutes in equilibration buffer (30% v/v glycerol, 2% w/v SDS, 50mM Tris, pH8.8, 0.02% bromophenol blue) containing 1% w/v DTT (reductive), and then for another 15 minutes in equilibration buffer containing 2.5% w/v Iodoacetamide (alkylating). In the Ettan Dalt Six system (GE Healthcare, USA), isoelectric focused proteins in the equilibrated IPG strips were separated in the second dimension using SDS PAGE. The proteins in the analytical gels were stained with Coomassie Brilliant Blue R Concentrate (Sigma-Aldrich, USA) after being fixed for ten hours in 50% methanol and 10% acetic acid. Melanie 9.0 2D gel analysis software was used to analyse protein spot detection, fold expression ratio, and spot patterns.

### In-Gel Trypsin Digestion

In-gel digestion and peptide extraction were performed according to the previously described protocol [17], with the experimental parameters described below. Protein spots of interest were manually removed from the 2D gels using a sterile scalpel and cut into approximately 1 mm³ pieces before transfer to microcentrifuge tubes. The gel pieces were subjected to destaining by incubation with 300 μL of 50 mM ammonium bicarbonate/acetonitrile (1:1, v/v) for 60 min with occasional vortexing. Following removal of the supernatant, the gel pieces were further treated with 500 μL acetonitrile at room temperature with intermittent vortexing until they became white and contracted. The solvent was then discarded, and the dried gel pieces were rehydrated with sequence-grade modified trypsin (13 ng/μL; Sigma-Aldrich, USA) prepared in 50 mM ammonium bicarbonate. Approximately 35 μL of the trypsin solution was used to cover the gel pieces, which were kept on ice for 45 min. Subsequently, 20 μL of 50 mM ammonium bicarbonate was added to maintain adequate hydration during digestion. The samples were incubated overnight at 37°C for enzymatic digestion. Peptides were subsequently recovered by adding 100 μL of extraction solution containing 5% formic acid/acetonitrile (1:2, v/v), followed by incubation at 37°C for 20 min with shaking. The resulting supernatant was transferred to a fresh tube and concentrated using a vacuum centrifuge. The dried peptide preparation was reconstituted in 20 μL of 0.1% formic acid, vortexed, and sonicated for 5 min, followed by centrifugation at 10,000 rpm for 12 min. The resulting supernatant was transferred to complete-recovery autosampler vials for subsequent LC/ESI-MS/MS analysis.

### Nanoflow Liquid Chromatography-tandem mass spectrometry

Protein identification was performed using nano-flow liquid chromatography coupled with electrospray ionization tandem mass spectrometry (nano-LC/ESI-MS/MS). Tryptic peptides were analyzed on a nanoACQUITY UPLC® system (Waters, Manchester, UK) operated using MassLynx 4.1 SCN781 software. Peptide separation was carried out by reversed-phase chromatography using 0.1% formic acid in water and acetonitrile as mobile phases A and B, respectively. Samples (3.0 μl) were introduced using a 5 μl partial-loop injection. Prior to analytical separation, peptides were desalted on a Symmetry® C18 trap column (180 μm × 20 mm, 5 μm; Waters) at 15 μl/min for 1 min and subsequently transferred to an HSS T3 C18 analytical column (75 μm × 200 mm, 1.8 μm; Waters). Separation was achieved using a 1–40% solvent B gradient over 43 min at 300 nl/min, followed by washing with 80% solvent B for 7 min and re-equilibration at 1% solvent B for 10 min. The column was maintained at 35°C. [Glu1]-Fibrinopeptide B human (Sigma) was used as the lock-mass calibrant ([M+2H]²⁺ = 785.8426), introduced through the reference sprayer at 300 nl/min. Each sample was analyzed in duplicate, with control injections performed between sample runs.

Peptide analysis was carried out on a SYNAPT® G2 High-Definition MS™ (HDMSE) system (Waters) equipped with a nano-electrospray ionization source. The instrument was operated in positive-ion mode with a capillary voltage of 3.5 kV, sample cone voltage of 50 V, extraction cone voltage of 6 V, and an IMS nitrogen gas flow of 90 mL/min. Ion mobility separation was performed using the T-Wave™ device with a pulse height of 40 V and a wave velocity of 800 m/s. During the IMS cycle, the travelling-wave amplitude was ramped from 8 to 20 V over 100% of the cycle. A NanoLockSpray™ source was used for mass calibration, with lock-mass measurements acquired at 45-s intervals. The TOF analyser was calibrated using 500 fmol/μL [Glu1]-Fibrinopeptide B human (Sigma-Aldrich, USA), allowing detection of ions in the 50–1200 m/z range. Data were acquired in resolution mode (V mode) with a resolving power of 18,000 FWHM and recorded in continuum format. HDMSE acquisition involved alternating between low- and elevated-energy functions. Function 1 generated low-energy MS spectra, whereas Function 2 acquired elevated collision-energy spectra combined with ion-mobility separation. For Function 2, the collision energy was maintained at 4 eV in the trap region and ramped from 20 to 45 eV in the transfer region to facilitate peptide fragmentation. Spectra were collected for 0.9 s per function with an interscan interval of 0.024 s.

### Mass Spectromtery Data analysis

The ion mobility-enhanced MSE data were processed using Progenesis QI for Proteomics version 3.0 (Non-Linear Dynamics, Waters) for protein identification. Following data acquisition, lock-mass correction was performed as part of the data-processing workflow. Noise thresholds for the low- and high-energy spectra were automatically determined by the ion-accounting function of the software. Protein identification was carried out by searching the curated human protein entries available in the UniProt database, with the false discovery rate (FDR) set to 1%. Protein assignments were accepted when the identification criteria included at least one matched fragment ion for a peptide, a minimum of three fragment-ion matches at the protein level, and at least one matched peptide per identified protein. Methionine oxidation was considered a variable modification, whereas carbamidomethylation of cysteine residues was specified as a fixed modification. Trypsin was selected as the digestion enzyme, allowing one missed cleavage.

### Functional enrichment, signaling pathways and protein-protein interactions analysis

Functional enrichment analysis of the identified proteins was performed using the Enrichr platform (https://maayanlab.cloud/Enrichr/) [18,19,20]. Enriched Gene Ontology (GO) terms, including biological processes, molecular functions, and cellular components, were identified based on enrichment confidence scores and functionally clustered to determine the biological significance of the target proteins. Pathway enrichment analysis of the differentially expressed proteins was further conducted using the Kyoto Encyclopedia of Genes and Genomes (KEGG) and Reactome datasets available in Enrichr. Data visualization and graphical plots were generated using the ggplot2 package in R software (version 4.6)[21]. Protein–protein interaction (PPI) network analysis of the identified proteins was performed using the STRING database (https://string-db.org/cgi/input.pl) [22] where proteins were clustered based on known and predicted interactions available in the database. Furthermore, physical interactions and gene co-expression patterns within the PPI network were analyzed using GeneMANIA (https://genemania.org/)[23].

### Gene expression analysis of differentially expressed proteins in SCF stimulated acute megakaryoblastic leukemia cells on TCGA-LAML patient samples

Gene expression analysis of stem cell factor (SCF)-activated proteins in acute megakaryoblastic leukemia cells was performed using the UALCAN platform (https://ualcan.path.uab.edu/)[24]. The TCGA-LAML cohort was selected, comprising 150 tumor samples from patients with acute myeloid leukemia (LAML). An interactive heatmap was generated to visualize the expression patterns of differentially expressed genes across the tumor samples. Furthermore, the expression profiles of these genes among different molecular and clinical subtypes of LAML were analyzed using the same platform. A comparative study of tumour and normal gene expression was conducted with gene expression data sourced from the GDC-TCGA-LAML dataset alongside normal samples from the GTEx database retrieved via UCSC Xena (https://xena.ucsc.edu/)[25]. A comprehensive study was conducted using 150 tumour specimens and 150 normal specimens. The outcomes were depicted as box plots created using the ggplot2 tool in R software (version 4.6), and statistical significance was determined using the log rank test in R.

### Receiver Operating Characteristic (ROC) and Co-expression analysis of LAML-genes

Diagnostic potential of leukemia-associated gene identified from stem cell factor (SCF)-activated acute megakaryoblastic leukemia cells was performed using the pROC package in R to assess their biomarker potential in acute myeloid leukemia (LAML). Receiver operating characteristic (ROC) curve analysis was conducted by plotting sensitivity against specificity, and the diagnostic performance of individual genes was evaluated based on the area under the curve (AUC) values. Genes exhibiting higher AUC values were considered to have greater diagnostic accuracy for LAML classification. Furthermore, co-expression analysis of the identified LAML gene signatures was performed using the GEPIA3 platform (http://gepia2.cancer-pku.cn/#index) [26] to investigate their association with known LAML-related genes. The analysis was used to evaluate the correlation patterns and potential functional relationships between SCF-activated gene signatures and established leukemia-associated genes.

### Survival and prognostic analysis of LAML associated genes in patients

Survival and prognostic analyses of stem cell factor (SCF)-activated leukemia-associated gene signatures were performed using survival data obtained from GDC-TCGA-LAML cohort, comprising 150 patients with acute myeloid leukemia (LAML), accessed through the UCSC Xena (https://xena.ucsc.edu/) platform. A univariable Cox regression model was subsequently constructed using the rms package in R to assess the combined prognostic performance of significant gene signatures and to estimate their contribution to survival outcomes in LAML patients. Representative hazard ratio and forest plots were generated using the forest model package in R to visualize the prognostic significance of candidate genes with corresponding 95% confidence intervals. To reduce overfitting and remove superfluous or anomalous variables, LASSO regression analysis was performed utilising the glmnet package in R, integrating transcriptome and survival data from TCGA-LAML patients. Additionally, Kaplan–Meier survival analysis was conducted utilising the survival and survminer packages in R to evaluate the overall survival outcomes between patients exhibiting high and low gene expression levels.

### Prognostic nomogram model for overall survival in LAML patients

Prognostic nomogram model was created using all significantly identified hazardous genes associated with overall survival in the TCGA-LAML cohort. The prognostic nomogram model was constructed using the rms package in R to predict 1, 3, and 5-year overall survival (OS) probabilities of LAML patients based on the expression profiles of selected prognostic genes signatures retained after LASSO filtering. The prognostic performance and reliability of the nomogram were further evaluated using calibration plots, time-dependent receiver operating characteristic (time-ROC) analysis, and decision curve analysis (DCA) implemented through the caret, timeROC, and dcurves packages in R, respectively.

### Quantitative Real Time PCR

Total RNA was extracted from both SCF-stimulated (100 ng/ml) and non-stimulated Mo7e cells utilising the RNeasy Mini Kit (Qiagen, Germany) in accordance with the manufacturer’s instructions. cDNA was generated via the RevertAid First Strand cDNA Synthesis Kit (Thermoscientific, USA) with Oligo (dT) primers in accordance with the manufacturer’s instructions. qRT-PCR was conducted on the Lightcycler 96 apparatus (Roche, Switzerland) utilising the KicqStart SYBR green Master mix from Sigma Aldrich, USA. Primers for quantitative reverse transcription polymerase chain reaction were acquired from Sigma Aldrich, USA. The expression of the target genes was standardised against the housekeeping gene GAPDH and shown as a relative fold change. P values were determined with an unpaired one-tailed Student’s t-test.

## Results

### 1. Global protein expression profiling in Stem Cell Factor stimulated human Acute Megakaryoblastic Leukemia cells

We attempted to understand the c-Kit signaling at proteomic level in acute megakaryoblastic leukemia cells using 2D gel electrophoresis. The cultured Mo7e cells (Fig S1) were either non-stimulated or stimulated with 100 ng/ml SCF for 24 hours (Fig S2). Differentially expressed proteins between non-stimulated and SCF stimulated Mo7e cells were studied using 2D gel electrophoresis and analyzed by Melanie 9.0 software (Fig 1). A total of 591 protein spots were observed of which 247 spots were up-regulated and 204 spots were down-regulated during SCF stimulation (Fig S3). The differences in protein spot intensity or size between non stimulated and SCF stimulated Mo7e cells were considered significant only when the fold change is greater than 1.5 and observed in two replicate gels obtained from two independent experiments (Fig S4). Significantly expressed protein spots, which are highly up- or down-regulated were excised and identified using MALDI-TOF/MS or LC-MS/MS. LC-MS/MS data was subjected for analysis by Progenesis QI software and proteins with highest confidence score and maximum number of unique peptides were selected for further validation and pathway analysis. The list of identified proteins with their accession numbers and confidence score is shown in Table 1. Furthermore, to confirm the protein spots identified by LC-MS/MS and MALDI-TOF/MS analysis, the theoretical molecular weight and isoelectric point (pI) of all the identified proteins were compared with the molecular weight and pI observed in 12% acrylamide gel. In all the conditions, the molecular weight and pI of the proteins in 12% acrylamide gel were consistent with the theoretical molecular weight and pI of the identified proteins.

**Fig. 1.**
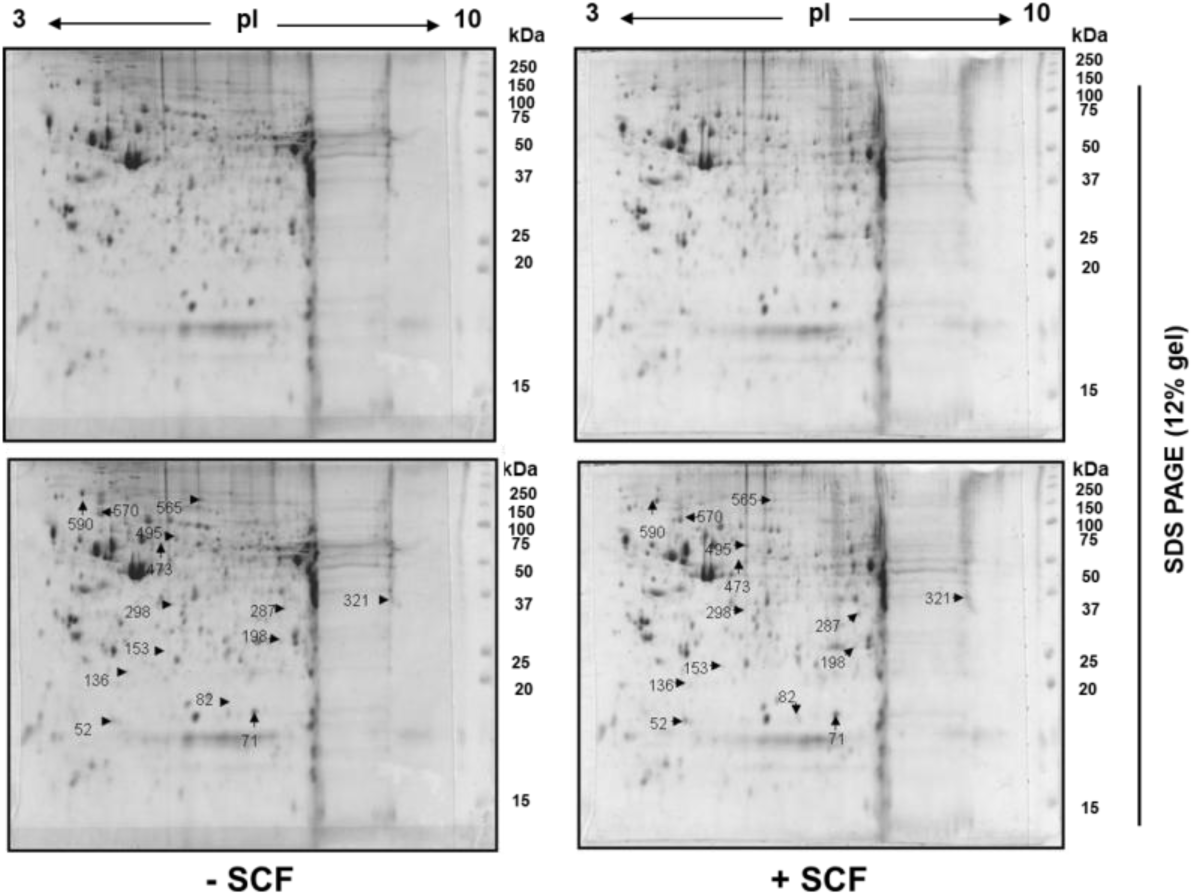
Two-dimensional (2D) electrophoresis profile of non-stimulated and SCF stimulated Acute Megakaryoblastic Leukemia cells (Mo7e). (A) Whole-cell lysates (900 μg) from both non-stimulated and SCF-stimulated Mo7e cells were fractionated by 2-D PAGE employing pH 3–10 IPG strips. The upper panel displays representative Coomassie-stained gels of both non-stimulated and SCF-stimulated Mo7e cells. (B) Quantitative analysis of protein spots by Melanie 9.0 two-dimensional gel analysis program. The figures in the gel denote markedly significant upregulated and downregulated protein locations chosen for identification with MALDI-TOF/MS or LC-MS/MS.

**Table-1:**
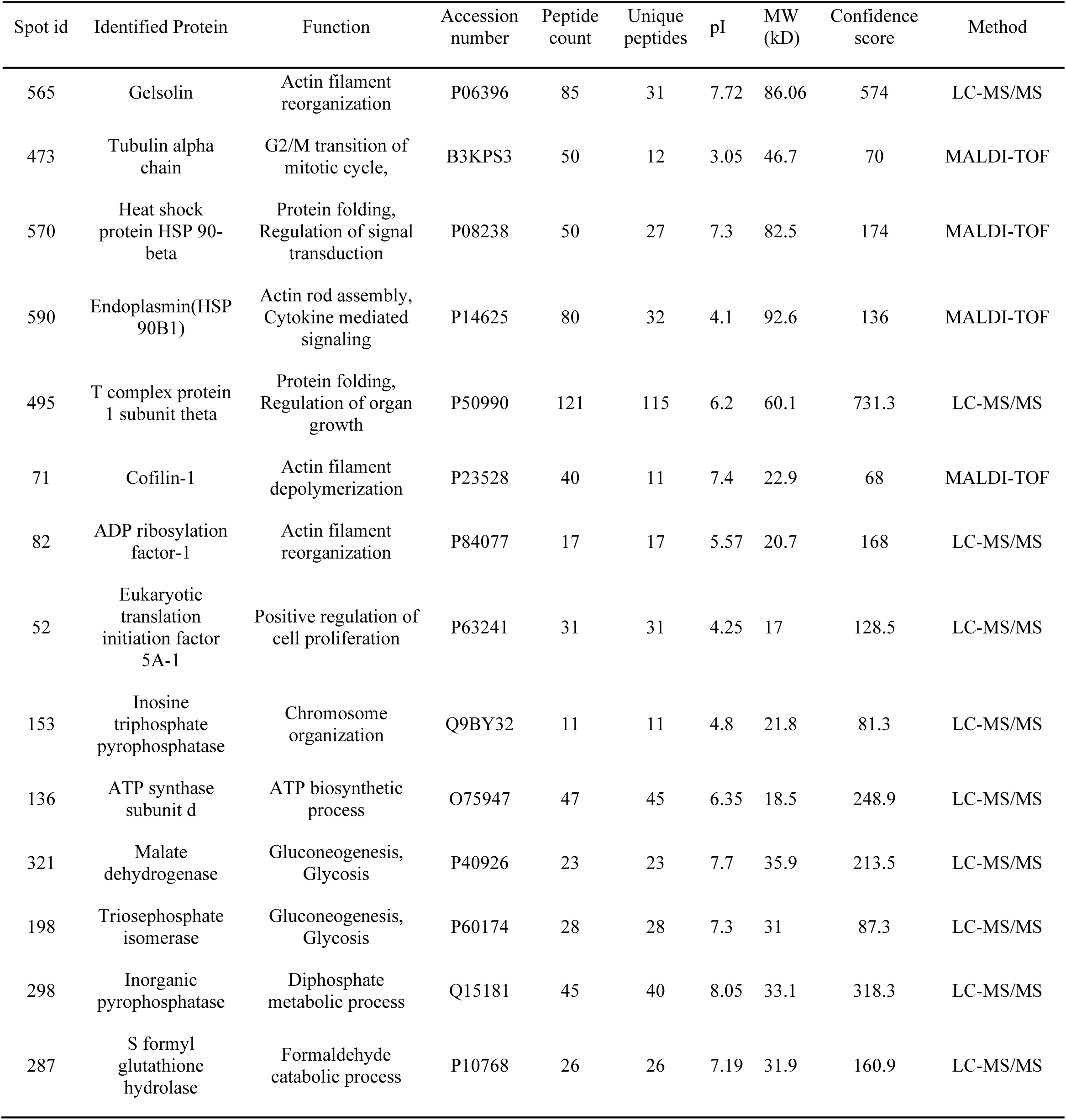
List of proteins identified in non-stimulated and SCF stimulated Acute megakaryoblastic leukemia cells.

In this study, we observed that cofilin-1 (CFL1), Endoplasmin (HSP90B1), T-complex protein 1 subunit theta (CCT8), eukaryotic translation initiation factor 5A1 (EIF5A), malate dehydrogenase, triosephosphate isomerase (MDH2), inorganic pyrophosphatase (PPA1), s-formyl glutathione hydrolase (ESD) were up-regulated and gelsolin (GSN), tubulin alpha (TUBA8), heat shock protein 90 beta (HSP90AB1), ADP ribosylation factor 1 (ARF1), inosine triphosphate pyrophospatase (ITPA) and ATP synthase subunit D (ATP5F1D) were down-regulated upon activation of c-Kit in human leukemic megakaryocytes.

### 2. Stem Cell Factor induces the expression of proteins involved in metabolism, cell cycle and actin reorganization

Gene ontology analysis of the identified proteins was performed using Enrichr to investigate Kit signaling regulated biological process, molecular function and cellular components in human leukemic megakaryocytes. Enrichr analysis revealed that the 14 identified proteins were enriched in biological process such as energy pathways, metabolism, endosome transport, ion transport, cell growth and maintenance, cell communication and signal transduction. Enrichr based molecular function analysis revealed that these proteins were involved in translation factor, chaperone, hydrolase, heat shock protein, isomerase, catalytic, GTPase and transporter activity. Further, the proteins were also found to be associated with structural constituent of cytoskeleton and cytoskeletal protein binding. Investigation on the distribution of identified proteins in the cell revealed that they were highly enriched in exosomes, cytoplasm, mitochondria, lysosome, cytosol and less enriched in plasma membrane, cytoskeleton, nucleolus, nucleus and centrosome (Fig. 2 A-C). Pathway enrichment analysis was conducted to investigate the biological activities and pathways linked to the discovered differentially expressed proteins. The examination disclosed that most proteins were chiefly engaged in protein metabolism, such as EIF5A, ARF1, GSN, MDH2, PPA1, and CCT8. Moreover, HSP90B1 and TUBA8 were identified as linked to cell cycle regulation, suggesting a potential role in the proliferation of leukemic cells. Furthermore, GSN was implicated in the regulation of the actin cytoskeleton, highlighting its potential involvement in cellular motility and structural organization, whereas HSP90B1 was enriched in the PI3K–AKT signaling pathway, a key pathway known to regulate cell survival, proliferation, and tumor progression (Fig.2 D). Overall, these findings suggest that the identified proteins may contribute to AML progression through coordinated regulation of metabolic processes, cell cycle progression, cytoskeletal remodeling, and oncogenic signaling pathways.

**Fig. 2.**
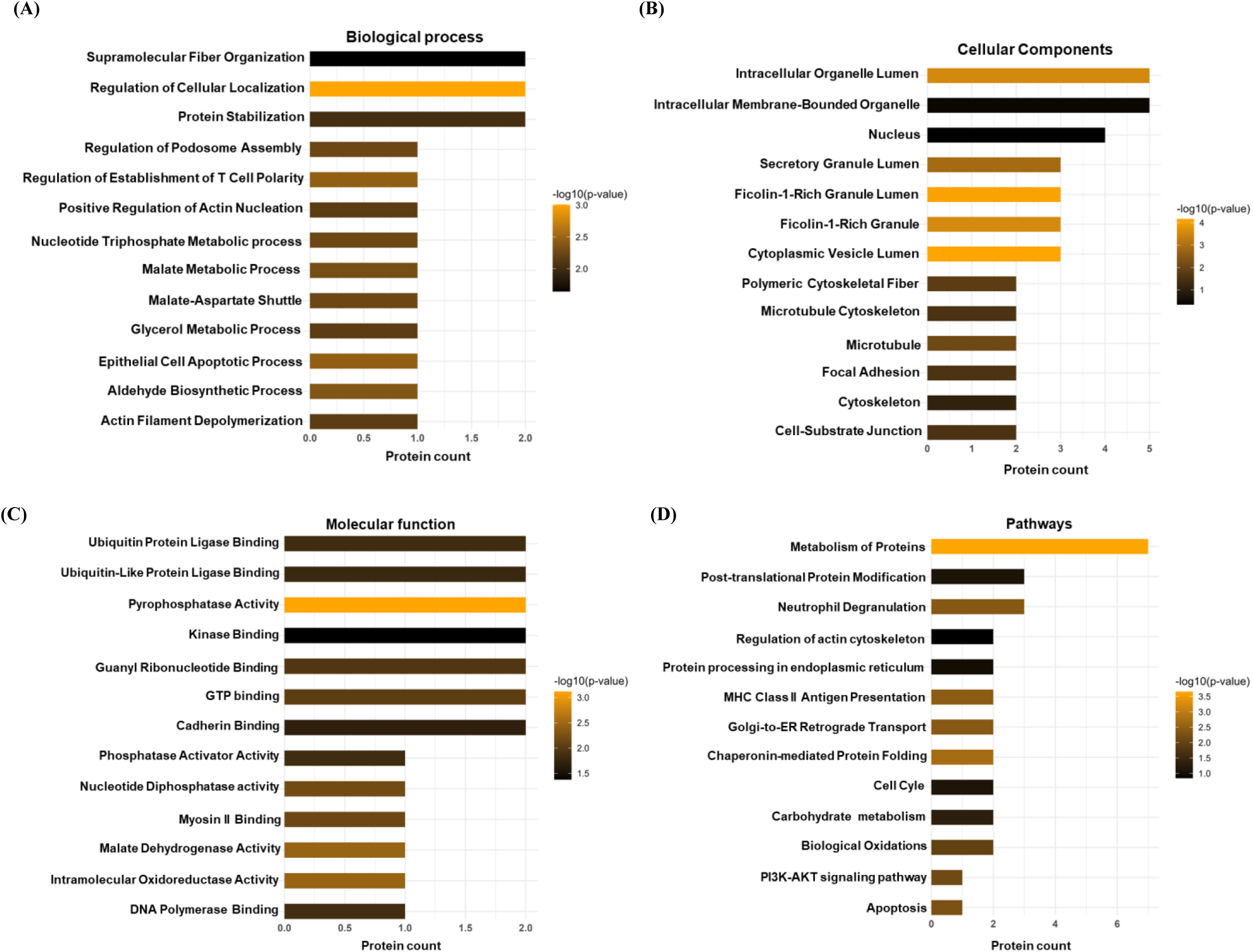
Functional enrichment analysis of differentially expressed proteins in SCF stimulated Acute Megakaryoblastic Leukemia cells. Enrichment analysis was performed using Enrichr, and the plots were generated using the ggplot2 package in R. Bar plot showing the top enriched (A) biological processes, (B) cellular components, (C) molecular functions and (D) signaling pathways.

### 3. Protein-protein interaction analysis reveals high physical and co-expression interaction among Stem Cell Factor induced proteins

To investigate the functional association among the identified proteins, a protein–protein interaction (PPI) network was constructed using the STRING database. The resulting interaction network demonstrated a significant enrichment of protein interactions (PPI enrichment *p* = 4.38 × 10⁻⁵), indicating that the identified proteins are biologically connected and functionally related rather than occurring as random associations (Fig. 3A). The network comprised 12 nodes and 17 edges, with an average node degree of 2.8 and an average local clustering coefficient of 0.73, suggesting a considerable level of interconnectivity and coordinated functional involvement among these proteins. Further we performed, gene– gene interaction analysis using GeneMANIA. The generated network revealed prominent physical interactions (49.8%) and co-expression patterns (26%) among the genes, highlighting their potential cooperative roles in shared biological processes (Fig. 3B). Additional interaction categories, including predicted and pathway-based associations, further supported the functional relatedness of these genes.

**Fig. 3.**
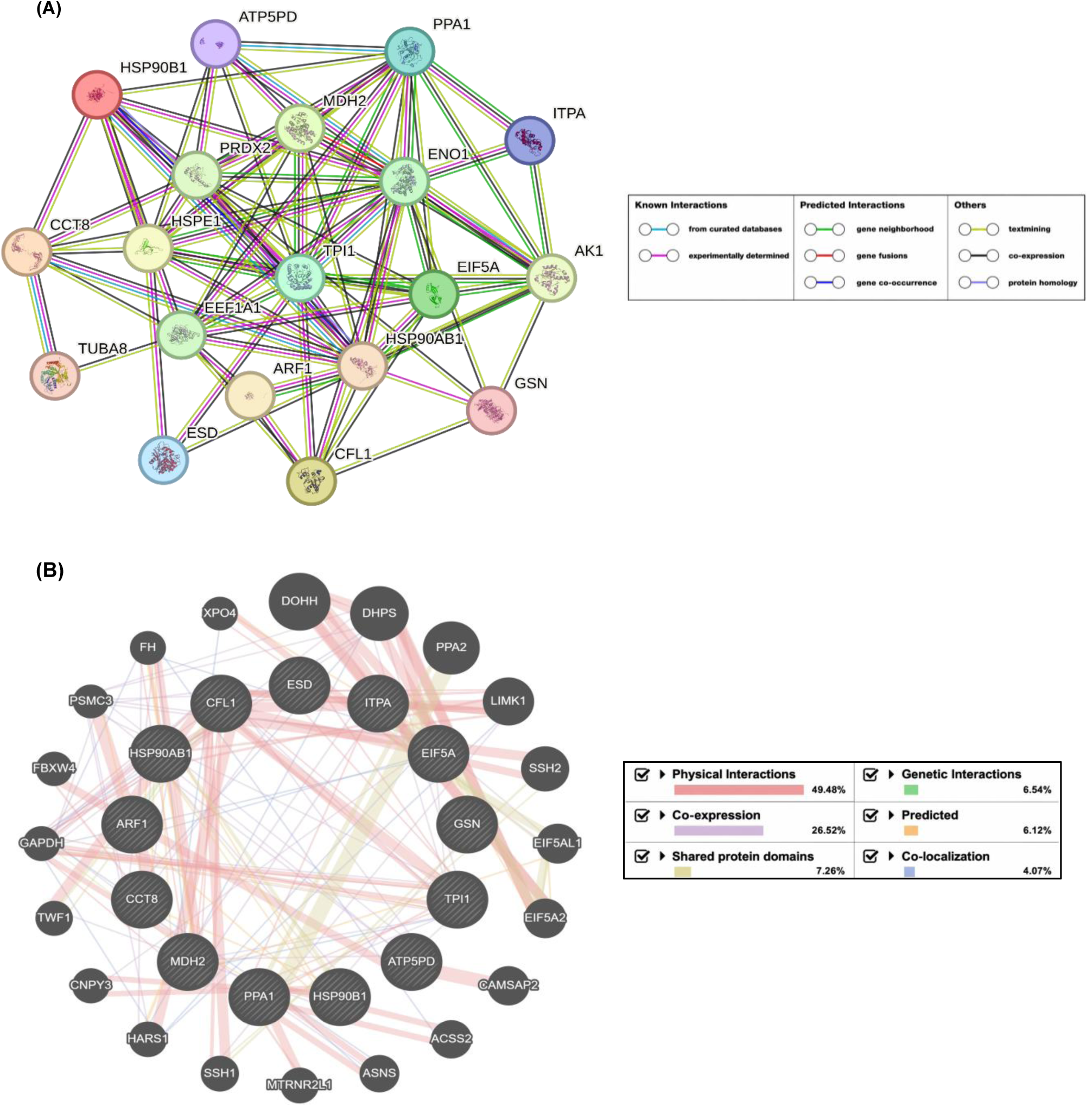
Protein-protein interaction network of identified proteins in SCF stimulated Acute Megakaryoblastic Leukemia cells. (A) STRING network: the nodes of the network (circles in the figure) represent proteins, while the edges of the network (the lines between the nodes) represent various interactions between the proteins. (B) Gene-Mania interaction network of identified proteins in terms of percentage physical interactions, Co-expression, Shared-protein domains, Genetic interactions and Co-localization.

### 4. Stem Cell Factor induced proteins in acute megakaryoblastic leukemia cells exhibits strong diagnostic and prognostic potential in Acute myeloid leukemia

To investigate the potential role of the identified differentially expressed proteins in acute myeloid leukemia (AML), the corresponding gene expression profiles were analyzed using the TCGA-LAML cohort obtained from the GDC portal and accessed through UCSC Xena. The LAML cohort consisted of gene expression data from 150 patient samples. Comparative analysis revealed that CFL1, EIF5A, ESD, HSP90B1, MDH2, GSN, and TPI1 exhibited elevated expression levels across AML patient samples. Notably, Kit, the receptor for Stem Cell Factor (SCF), was also found to be highly expressed in the cohort, suggesting a potential association between SCF-mediated signaling and the observed expression pattern of these genes (Fig 4A). Furthermore, expression analysis across different AML subtypes demonstrated that these genes were consistently expressed irrespective of subtype classification (Fig S5).

**Fig. 4.**
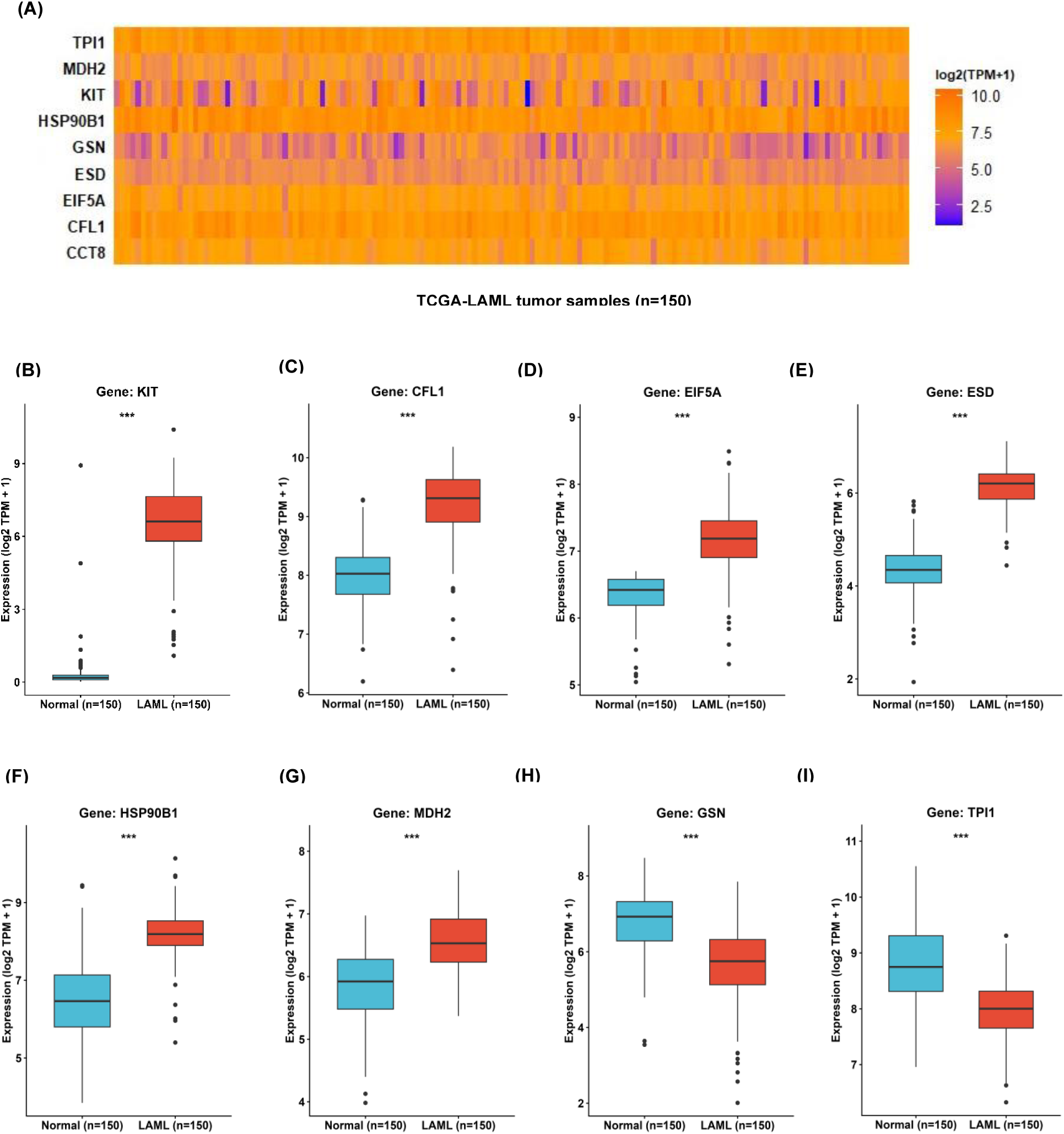
Gene expression analysis of differentially expressed proteins in SCF stimulated Acute Megakaryoblastic Leukemia cells in TCGA-LAML patient samples. Gene expression data for LAML patient samples were obtained from UCSC Xena, and plots were generated using the ggplot2 package in R. (A) Heat map showing the expression pattern of differentially expressed proteins. (B–I) Box plots showing the relative gene expression of differentially expressed proteins in LAML tumor and normal samples. Normal whole blood expression data were obtained from GTEx for comparison. Statistical significance between groups is indicated as follows: *p< 0.05, **p< 0.005, ***p< 0.0005.

To further evaluate differential gene expression, tumor-versus-normal comparison was performed using whole blood transcriptomic data obtained from the GTEx Portal as a normal control dataset. The analysis demonstrated that KIT, CFL1, EIF5A, HSP90B1, and MDH2 were significantly overexpressed in AML samples compared to normal blood tissues, whereas GSN and TPI1 showed reduced expression in AML patient samples (Fig 4 B-I). Further we validated these gene expression using extermenal AML data set GSE15061(25). Based on these observations, it may be speculated that SCF-mediated activation of c-Kit signaling could contribute to the dysregulated expression of downstream genes involved in leukemogenesis. To assess the diagnostic potential of these genes, receiver operating characteristic (ROC) curve analysis was performed. Most candidate genes demonstrated strong discriminatory performance, with area under the curve (AUC) values exceeding 0.5, while CFL1, GSN, CCT8, HSP90B1, and TPI1 exhibited comparatively higher AUC values, indicating promising diagnostic utility in AML (Fig 5A). Additionally, co-expression analysis revealed significant positive correlations between the identified gene signature and previously established AML-associated genes, further supporting their biological relevance in leukemia progression (Fig S6B).

**Fig. 5.**
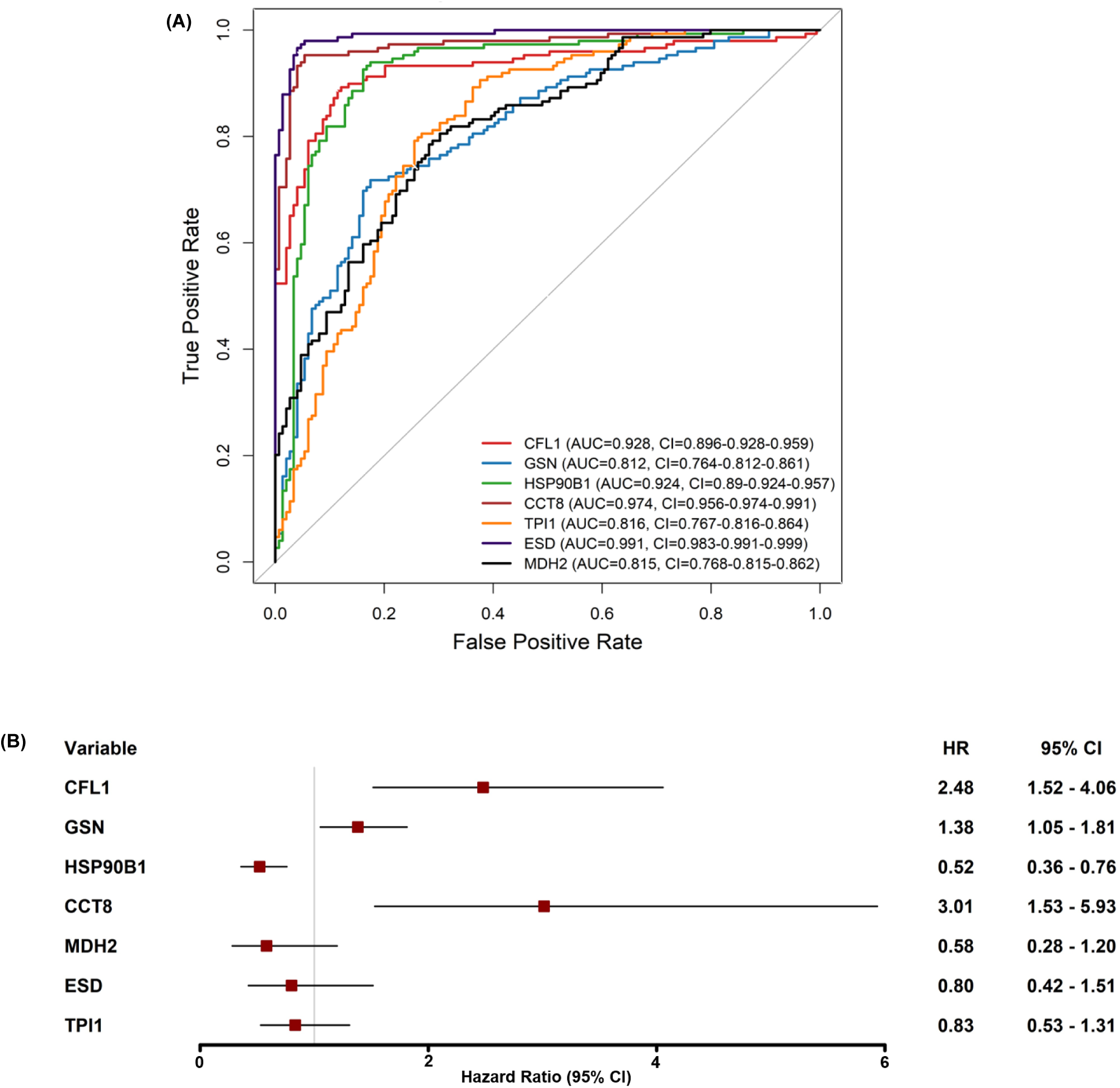
Receiver Operating Characteristic (ROC) and Univariable hazard analysis of LAML-genes in SCF stimulated Acute Megakaryoblastic Leukemia cells. (A) ROC curves for LAML gene signatures (CFL1, GSN, HSP90B1, CCT8, TPI1, ESD, and MDH2). The diagnostic performance is represented by the area under the curve (AUC) with 95% confidence intervals (CI). (B) Univariable Cox proportional hazards model showing hazard ratios (HRs) with corresponding 95% confidence intervals (CI) for gene signatures in LAML patient samples. HR > 1 indicates that higher gene expression is associated with poorer survival, HR = 1 indicates no significant association, and HR < 1 indicates a protective effect on patient survival.

### 5. Construction of prognostic nomogram model of survival for identified proteins

To determine the prognostic significance of the identified genes, univariate Cox proportional hazards regression analysis was performed using clinical survival data from the GDC-TCGA-LAML cohort. The analysis identified CFL1, GSN and CCT8 genes associated with elevated hazard risk, suggesting their potential involvement in poor patient prognosis (Fig 5b). Further, least absolute shrinkage and selection operator (LASSO) regression analysis was performed to eliminate redundant variables and identify the most robust hazardous genes. The coefficient path plot indicated deviation of TPI1, ESD, MDH2 and HSP90B1, leading to its exclusion from subsequent analyses, while CFL1 and GSN were retained and it confirms the hazardous potential of these gens (Fig S7). Subsequently, Kaplan–Meier survival analysis was conducted by stratifying patients into high- and low-expression groups. The results demonstrated that elevated expression of CFL1 (Fig 6A-B), GSN (Fig 6 C-D), and CCT8 (Fig 6 E-F) was significantly associated with reduced overall survival in AML patients, highlighting their prognostic relevance. To establish a prognostic prediction model, a nomogram for AML patient survival was constructed. The resulting nomogram model was designed to predict 5-year overall survival probability in AML patients. The predictive performance and reliability of the nomogram were further evaluated through calibration analysis, time-dependent ROC curves, and decision curve analysis (DCA), all of which demonstrated favorable predictive accuracy and clinical applicability (Fig. 7,8). The diagnostic performance of these nomograms was also validated in independently in AML data set GSE37642 (26). Collectively, these findings support the reliability of the proposed prognostic model and suggest that CFL1, GSN and CCT8 may serve as promising prognostic biomarkers for AML.

**Fig. 6.**
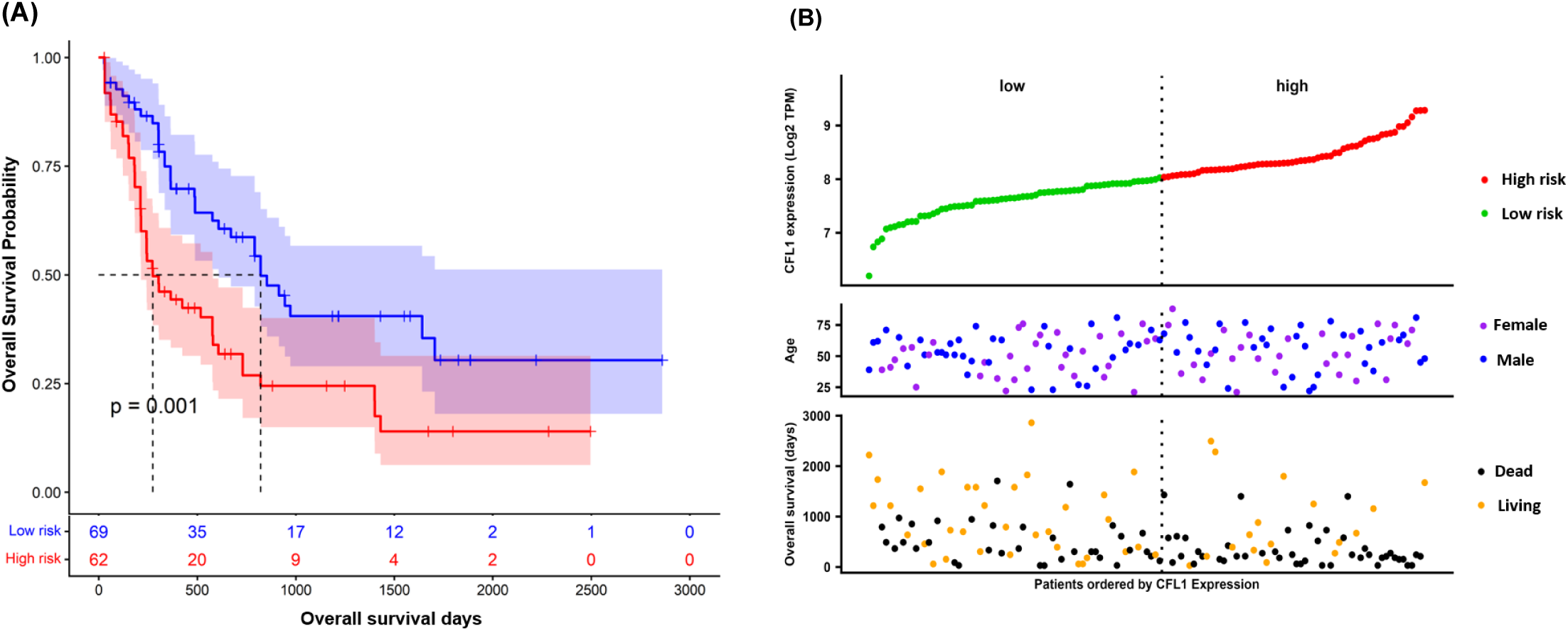
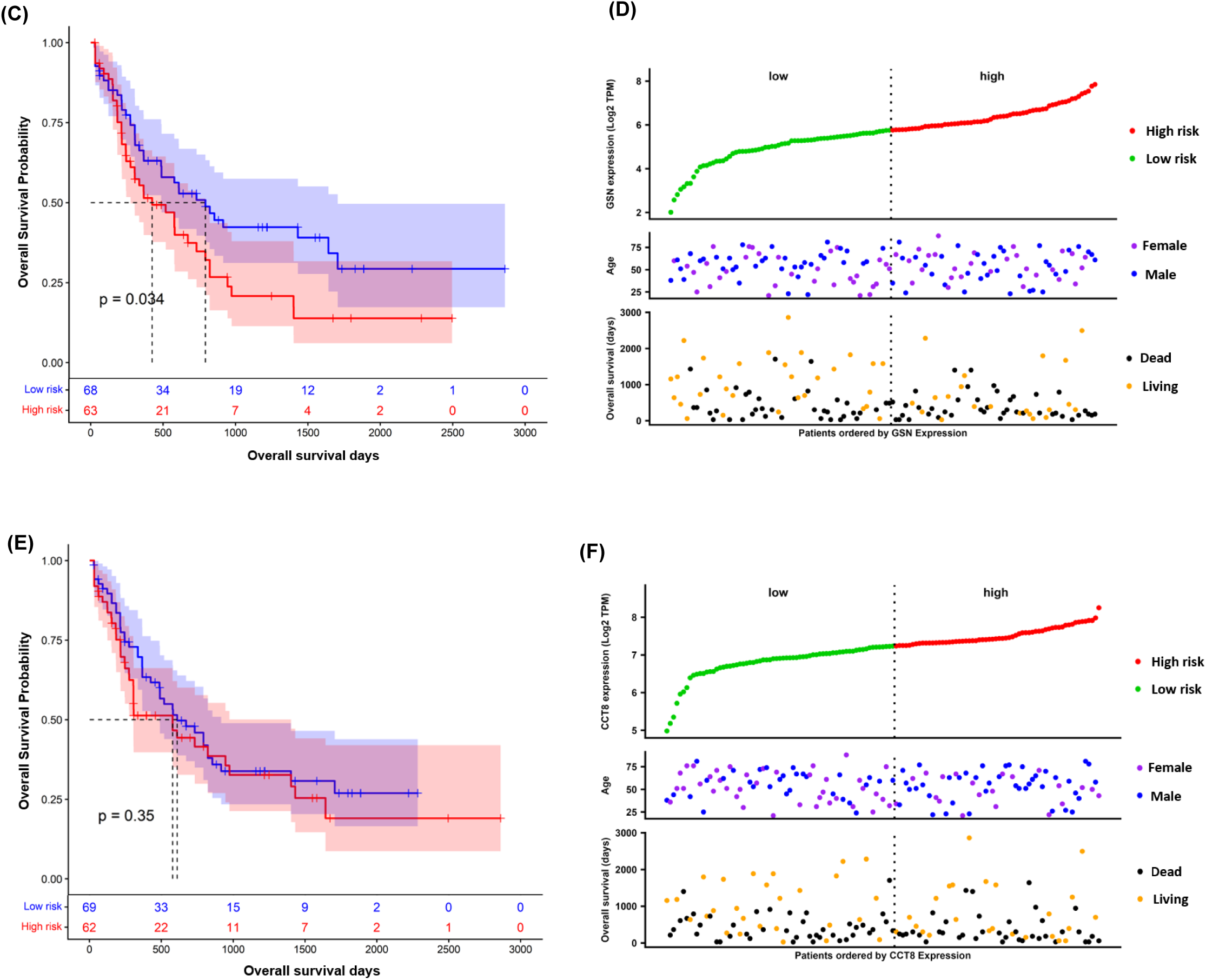
Survival and prognosis analysis of SCF stimulated LAML related genes in Acute Megakaryoblastic Leukemia cells. Survival analysis was used to assess the prognostic relevance of deleterious genes (CFL1, GSN, and CCT8). Kaplan–Meier survival curves (A-B) contrasting overall survival between groups with high and low expression of the prognostic gene CFL1. Kaplan-Meier survival curves (B-C) contrasting overall survival rates between high-expression and low-expression cohorts for the prognostic gene GSN. Kaplan–Meier survival curves (D-E) contrasting overall survival between groups with high and low expression levels of the prognostic gene CCT8. All statistical significance was evaluated via the log-rank test, with 95% confidence intervals shown.

**Fig. 7.**
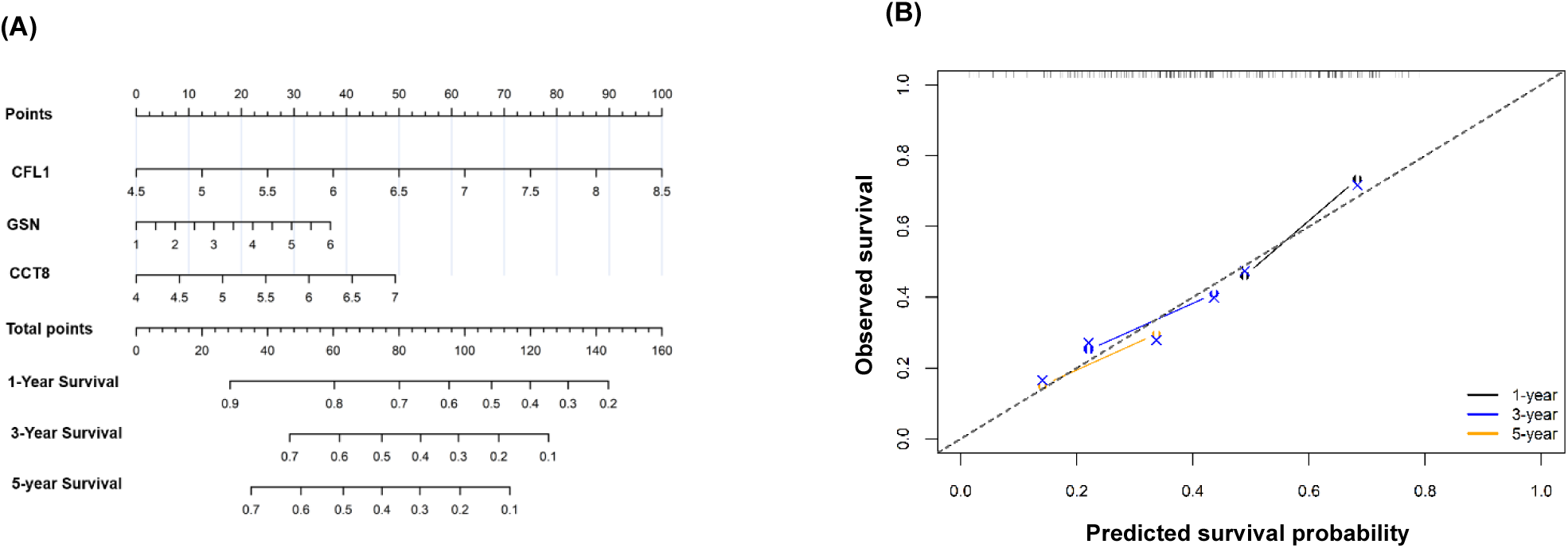
Construction of prognostic nomogram model of overall survival in Acute Myeloid Leukemia patients. (A) Prognostic nomogram was developed based on a univariable Cox proportional hazards regression model integrating three gene expression signatures (CFL1, GSN and CCT8) Independent prognostic factors were identified and incorporated into the model, and their corresponding regression coefficients were proportionally transformed into a point-based scoring system. Each variable is represented on an individual axis, where a vertical projection to the “Points” scale assigns a weighted score. The cumulative score (Total Points) is obtained by summing individual contributions and is subsequently mapped to predict 1, 3, and 5-year OS probabilities. (B) Calibration of the prognostic nomogram for overall survival. Calibration plots for 1, 3, and 5-year overall survival comparing predicted probabilities with observed outcomes. The dashed line represents perfect agreement. Solid lines denote bootstrap bias-corrected estimates (B = 1000), with points derived from groups of patients. The close alignment with the reference line indicates good calibration of the model.

**Fig. 8.**
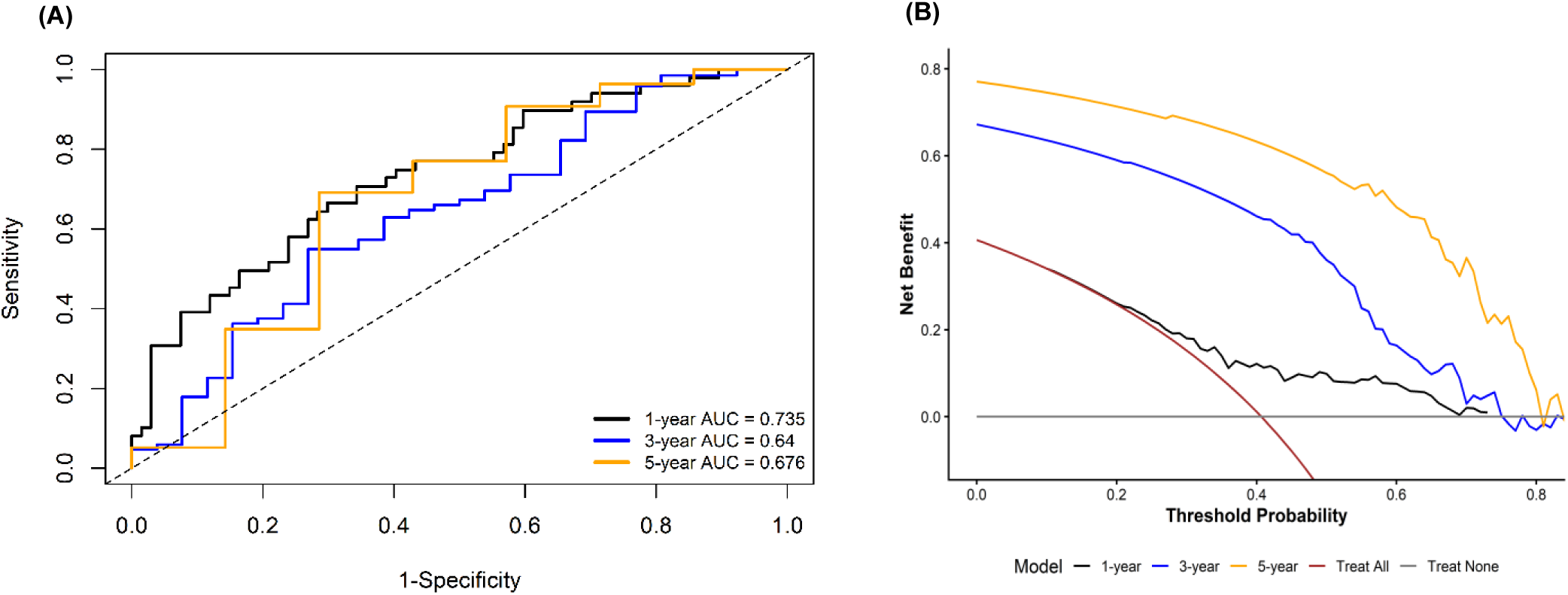
Validation of prognostic nomogram model. (A) Time ROC curves of the prognostic nomogram for overall survival. Time receiver operating characteristic (ROC) curves assessing the prognostic efficacy of the nomogram for overall survival at 1, 3, and 5 years. The relevant areas under the curve (AUCs) were 0.73, 0.64, and 0.67. The diagonal dashed line signifies arbitrary forecasting. The model has a reasonable capacity for discrimination at various time intervals. (B) Evaluation of the decision curve for the prognostic nomogram for overall survival. Decision curve analysis (DCA) illustrating the standardised net advantage of the nomogram for forecasting overall survival at 1, 3, and 5 years across various threshold probabilities. The black, blue, and orange lines denote the net advantage for predictions spanning 1 year, 3 years, and 5 years, respectively. The grey lines denote the benchmark techniques for treating all patients (“All”) or treating no patients (“None”). The nomogram exhibits greater net benefit across a wide spectrum of threshold probabilities when compared to standard techniques, suggesting its possible therapeutic use.

### 6. Cofilin-1 (CFL1), Gelsolin (GSN) and T-Complex protein-1 (CCT8) in stem cell factor stimulated acute megakaryoblastic leukemia cells are promising prognostic biomarkers for AML

The differential expressions of predicted biomarkers were further validated at the transcript level using quantitative real time – PCR. The identified proteins were picked and studied for their mRNA expression levels. The mRNA expression levels of Cofilin-1, Gelsolin and T-complex protein 1 subunit theta along with Endoplasmin, were analyzed in 12 and 24 hours of SCF stimulated Mo7e cells. Analysis of mRNA expression levels revealed that Endoplasmin, T-complex protein 1 subunit theta, and Cofilin were up-regulated and Gelsolin was down-regulated in SCF stimulated Mo7e cells (Fig 9). The fold expression of mRNA levels was similar to that of protein, which further validates potential of the identified proteins by LC-MS/MS and MALDI-TOF/MS analysis as prognostic biomarkers for AML.

**Fig. 9.**
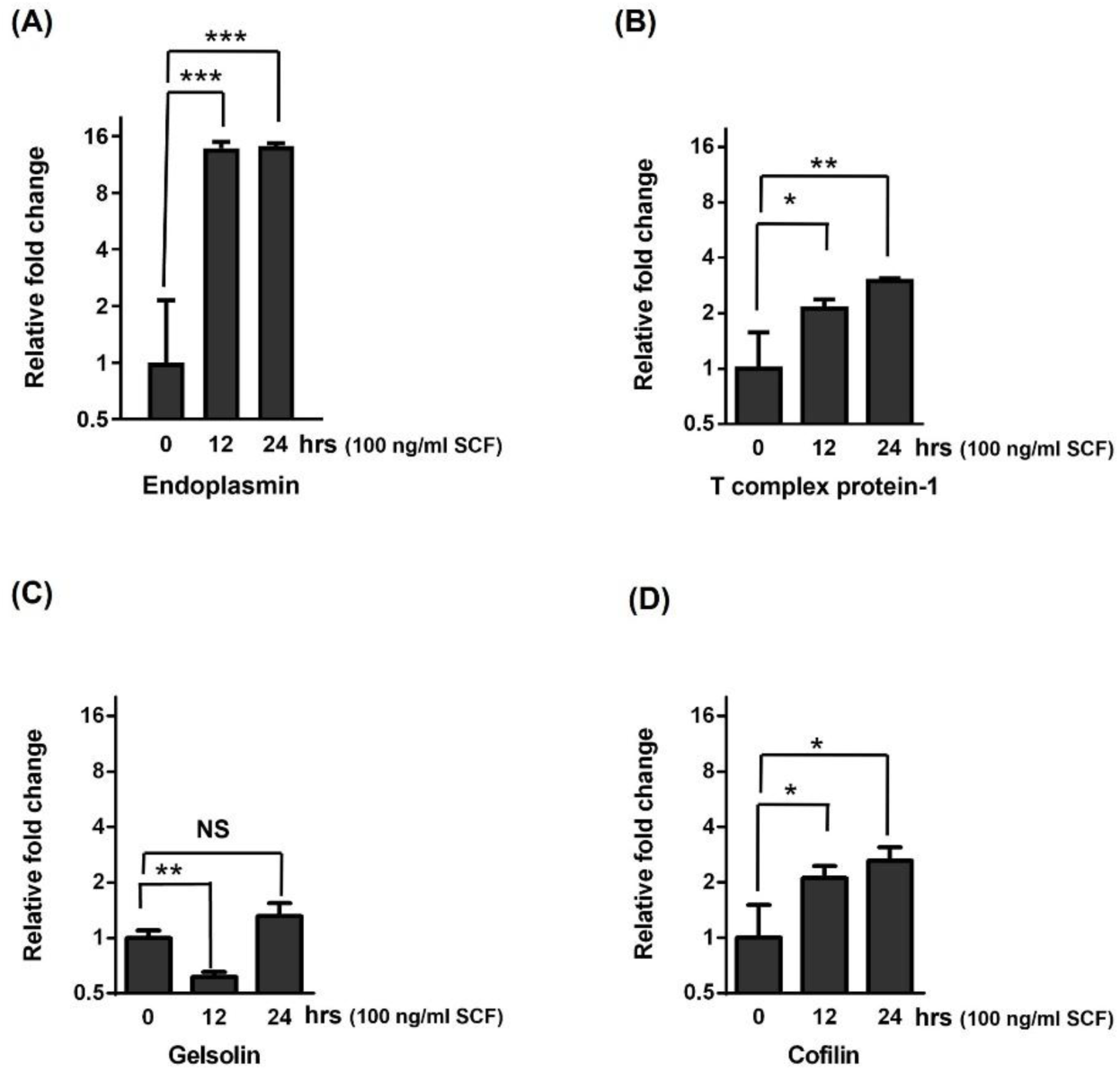
Quantitative real time PCR based validation of differentially expressed genes in SCF stimulated Acute Megakaryoblastic Leukemia cells (Mo7e). Expression levels of Kit signalling dependent genes were analysed by qRT-PCR after stimulating Mo7e cells with 100 ng/ml SCF for 0, 12 and 24 hours. Relative gene expression was normalized using GAPDH. Data are represented as mean ± SD. (n=3). p value is calculated using unpaired student t test. *p< 0.05, **p< 0.005, ***p< 0.0005.

## Discussion

The stem cell factor receptor or c-Kit, is a versatile type-3 receptor tyrosine kinase that is expressed in a various cancer type. Activation of c-Kit by its ligand, stem cell factor (SCF), has been shown to contribute to leukemogenesis by promoting the expression of proteins involved in leukemia cell proliferation, survival, migration, and resistance to apoptosis [27–30]. In the present study, we utilized SCF-responsive acute megakaryoblastic leukemia (M07e) cells as a model to investigate the proteomic changes following c-Kit activation. Through comparative proteomic approach (2D-GE-LC-MS/MS), we have identified 14 proteins exhibiting differential expression upon SCF treatment. Functional enrichment analysis revealed that these proteins are involved in the regulation of cytoskeletal organization (CFL1, GSN, TUBA8), protein folding (HSP90B1, HSP90AB1, CCT8), carbohydrate metabolism (MDH2, TPI1, PPA1), vesicular trafficking (ARF1), translational regulation (EIF5A), and nucleotide metabolism (ITPA). Additionally, signaling pathway analysis indicated that these proteins participate in pathways associated with protein metabolism, cytoskeletal dynamics and energy production. These observations are relevant, because metabolic reprogramming, increased protein synthesis, altered proteostasis, and actin remodeling have emerged as major process supporting leukemic cell proliferation, survival, and resistance to therapy [30–34]. Taken together, these findings suggest that SCF stimulation may induce a comprehensive metabolic shift in M07e cells via c-Kit signaling pathways. Furthermore, the significant physical interactions and co-expression patterns observed among these proteins imply a coordinated mechanism that underlies the cellular effects induced by SCF.

To understand the clinical relevance of these proteins, we examined their relative gene expression in TCGA-LAML patient dataset as well as independent GEO datasets. This approach constitutes a major strength of the present study, as proteomic findings are often constrained by the lack of validation in patient samples. Our analysis revealed that CFL1, EIF5A, HSP90B1, and MDH2 were consistently overexpressed in acute myeloid leukemia (AML), while GSN and TPI1 were significantly downregulated across both the TCGA and GEO datasets, in agreement with our proteomic findings. These reproducible expression patterns suggest that the identified proteins are not restricted to cell lines but instead represent conserved downstream effectors of SCF/c-Kit signaling in AML. Furthermore, receiver operating characteristic (ROC) and co-expression analyses demonstrated that all of these genes exhibit satisfactory diagnostic performance, underscoring their potential as candidate biomarkers for AML.

The prognostic potential of the genes was further explored, through multiple statistical approaches, including Cox regression hazard models, Random Forest and LASSO analyses. We found CFL1, CCT8 and GSN as top prognostic candidates. Kaplan–Meier survival analysis further confirmed their significant association with poor prognosis, indicating that SCF responsive genes have direct clinical relevance. Cofilin-1 (CFL1), a pivotal regulator of actin filament turnover, orchestrates cytoskeletal dynamics essential for leukemic cell proliferation, migration, and interaction with the bone marrow microenvironment [35–36]. Previous studies have identified CFL1 as a prognostic marker in various malignancies, including lung, breast, skin, and prostate cancers, where elevated expression correlates with poor outcomes and increased drug resistance [37–40]. In line with these findings, our study observed consistent upregulation of CFL1 following SCF stimulation in Mo7e cells and high expression in AML patients, establishing CFL1 as the most potent adverse prognostic indicator across both discovery and validation cohorts. Collectively, these results suggest that CFL1 may function as a pan-cancer prognostic marker warranting further investigation.

Gelsolin (GSN), another key modulator of actin cytoskeleton organization, exhibited significant downregulation after SCF stimulation and in Mo7e cells and in AML patient samples. Reduced GSN expression was generally associated with poorer overall survival, and hypogelsolinemia is frequently observed in AML during cancer progression [41]. Similar trends have been reported in gastric cancer patients [42]. Thus, GSN may serve as a valuable diagnostic marker, as its decreased expression is linked to unfavorable prognosis. T-complex protein (CCT8), is a subunit of the TRiC/CCT chaperonin complex involved in the folding of actin, tubulin, and various cell-cycle proteins, was markedly upregulated following SCF treatment and in AML specimens. Its elevated expression has also been documented in B-cell non-Hodgkin lymphoma, where it is associated with adverse outcomes, and knockdown of CCT8 in lung adenocarcinoma cells impairs proliferation and migration [43–44]. In addition to the proteomics data, we validated the gene expression of prognostic genes using qRT-PCR in SCF stimulated Mo7e cells. We observed upregulation of CFL1 and CCT8, along with the downregulation of GSN upon SCF stimulation, demonstrated a strong concordance between protein levels and transcript abundance. This indicates that the molecular changes reflect genuine biological responses to c-Kit activation rather than technical artifacts. Furthermore, the consistency observed between in vitro data and patient-derived transcriptomic datasets underscores the biological significance of the proposed biomarkers.

To translate these findings into clinical practice, we developed a prognostic nomogram based on CFL1, CCT8, and GSN expression. This model accurately predicted 1-, 3-, and 5-year overall survival, demonstrated reliable calibration, and exhibited strong time-dependent ROC performance and decision curve analysis, indicating potential clinical utility. Compared to larger gene signatures, this concise three-gene panel may offer greater practicality for routine clinical implementation. The model’s robustness was further confirmed using an independent external GEO cohort.

## Limitations

The proteomic analysis was performed using the Mo7e cells, which is although widely accepted as an experimental model for investigating SCF/c-Kit signaling, may not fully capture the biological heterogeneity of primary AML. So, it required further validation in primary leukemia cells. In addition, the present study focused primarily on identifying highly differentially expressing proteins, so total 14 proteins only we considered, however, in order to get mechanistic view a greater number of proteins need to identify. Functional studies involving gene knockdown, gene knock in, and pharmacological inhibition studies will be required to determine the precise contribution of individual proteins, particularly CFL1, GSN, and CCT8, to leukemic cell proliferation, apoptosis, migration, and therapeutic responses.

## Conclusions

In conclusion, the present study provides the first time global proteomic characterization of SCF-mediated c-Kit signaling in human megakaryoblastic leukemia cells and demonstrates that c-Kit activation induces coordinated regulation of proteins involved in cytoskeletal remodeling, protein homeostasis, metabolism, intracellular trafficking, and oncogenic signaling. Integration of experimental proteomics with TCGA-LAML transcriptomic validation, network analysis, diagnostic evaluation, survival analysis, and prognostic modeling identified CFL1, GSN, and CCT8 as clinically relevant downstream effectors of SCF/c-Kit signaling with significant diagnostic and prognostic potential in AML. These findings improve our understanding of the molecular mechanisms linking SCF/c-Kit signaling to leukemogenesis and establish a foundation for future mechanistic studies aimed at developing novel biomarkers and therapeutic targets for AML.

## Authors Contributions

A.K.R, G.G & S.A performed research, analyzed data and wrote the paper. A.S performed research, analyzed data. M.M designed the experiments, analyzed data and wrote the paper.

## List of Abbreviations

AKT: Protein Kinase B (PKB)
ARF1: ADP-Ribosylation Factor 1
CCND1: Cyclin D1
CXCR4: C-X-C Motif Chemokine Receptor 4
EIF5A: Eukaryotic Translation Initiation Factor 5A
ERK: Extracellular Signal-Regulated Kinase
ESD: S-formylglutathione hydrolase
GEO: Gene Expression Omnibus
GEPIA3: Gene Expression Profiling Interactive Analysis 3
GTEx: Genotype-Tissue Expression
HOXA9: Homeobox A9
HSP90B1: Heat Shock Protein 90 Beta Family Member 1
JAK1: Janus Kinase 1
LASSO: Least Absolute Shrinkage and Selection Operator
LC: MS/MS-Liquid Chromatography–Tandem Mass Spectrometry
MALDI: TOF-Matrix-Assisted Laser Desorption/Ionization–Time of Flight
MAPK: Mitogen Activated Protein Kinase
MDH2: Malate Dehydrogenase 2
MEIS1: Meis Homeobox 1
MYC: MYC Proto-Oncogene, BHLH Transcription Factor
PI3K: Phosphoinositide 3-Kinase
PPA1: Inorganic Pyrophosphatase 1
ROC: Receiver Operating Characteristics
RPMI: Roswell Park Memorial Institute Medium
RUNX1: Runt-Related Transcription Factor 1
STAT1: Signal Transducer and Activator of Transcription 1
STAT5: Signal Transducer and Activator of Transcription 5
STRING: Search Tool for the Retrieval of Interacting Genes/Proteins
TCGA: LAML-The Cancer Genome Atlas – Acute Myeloid Leukemia
TPI1: Triosephosphate Isomerase 1
TUBA8: Tubulin Alpha 8
UCLAN: University of Central Lancashire
UCSC: University of California, Santa Cruz

## Ethics approval and consent to participate

Not applicable

## Human and animal rights

Not applicable

## Consent for publication

Not applicable

## Availability of data and materials

The data sets used for this study in publicly available in TCGA, GTEx and GEO databases.

## Acknowledgements

This work was supported by ICMR (2020-3375). We thank Pondicherry University for providing Ph.D fellowship to Gopika Gopan and Sudhakar Arumugam. We also acknowledge DBT for providing fellowship to Anand Kuttanparambil Ravi. We tank Proteomics Core facility, RGCB, Trivandrum, Kerala for LC-MS/MS and MALDI-TOF analysis. We are grateful to DST-FIST, Government of India [SR/FST/LS-I/2019/426(C)] for infrastructure facilities in Department of Microbiology.

## Supplementary materials

Additional information associated with this article including tables and images are available in the supplementary materials.

