## Supplementary materials for "Global protein expression profiling in stem cell factor stimulated human Acute megakaryoblastic leukemia cells identifies CFL1, GSN and CCT8 as prognostic biomarkers for Acute Myeloid Leukemia"

### Supplementary data

**Table S1: Primers used in qRT-PCR analysis**

| Gene/Protein | Forward Primer 5'-3' | Reverse Primer 3'-5' |
|---|---|---|
| GAPDH | TCACCAGGGCTGCTTTTAAC | TGACGGTGCCATGGAATTTG |
| HSP90B1 (Endoplasmin) | TTCAAAGGAAAGTGATGACC | GCATCATATCATGGAAGTCG |
| CCT8 (t complex protein subunit theta) | CTTCCTAGATTGACACCTCC | TACTGCCCTTTCTATGTCATC |
| GSN (Gelsolin) | ACAGATCTGGAGAATCGAAG | TATAGATTATCTGCCCCTGG |
| CFL1 (Cofilin) | CTATGATGCAACCTATGAGAC | CTGTCAGCTTCTTCTTCTTGATG |

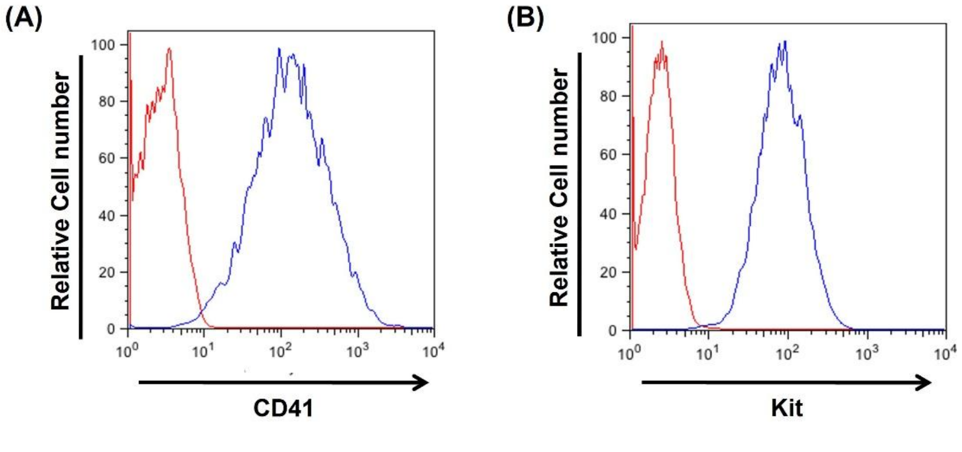


**Fig. (S1). Surface expression levels of Kit and CD41 in Acute Megakaryoblastic Leukemia cells**. Cells were analyzed for Kit and CD41 surface expression. The blue color represents antibodies against Kit or CD41 and red color represent the corresponding isotype control.

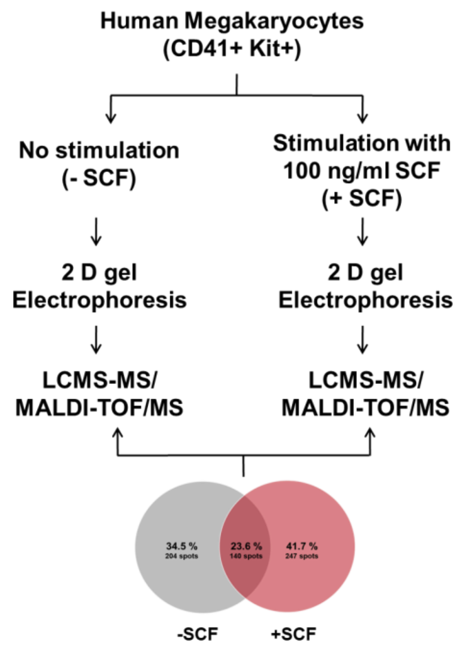


**Fig. (S2). Schematic representation of experimental workflow in non-stimulated and SCF stimulated Acute Megakaryoblastic Leukemia cells.**

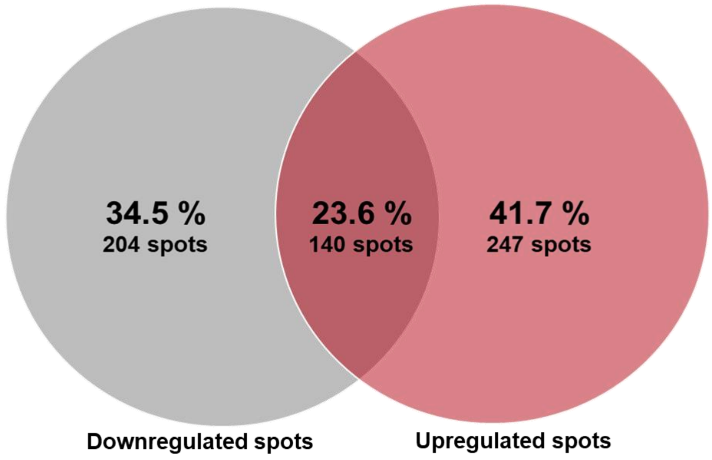


**Fig. (S3). Comparison of protein expression in non-stimulated and SCF stimulated Acute Megakaryoblastic Leukemia cells.** Venn diagram represents the percentage and number of differentially expressed proteins in non-stimulated and KL stimulated Acute Megakaryoblastic Leukemia cells.

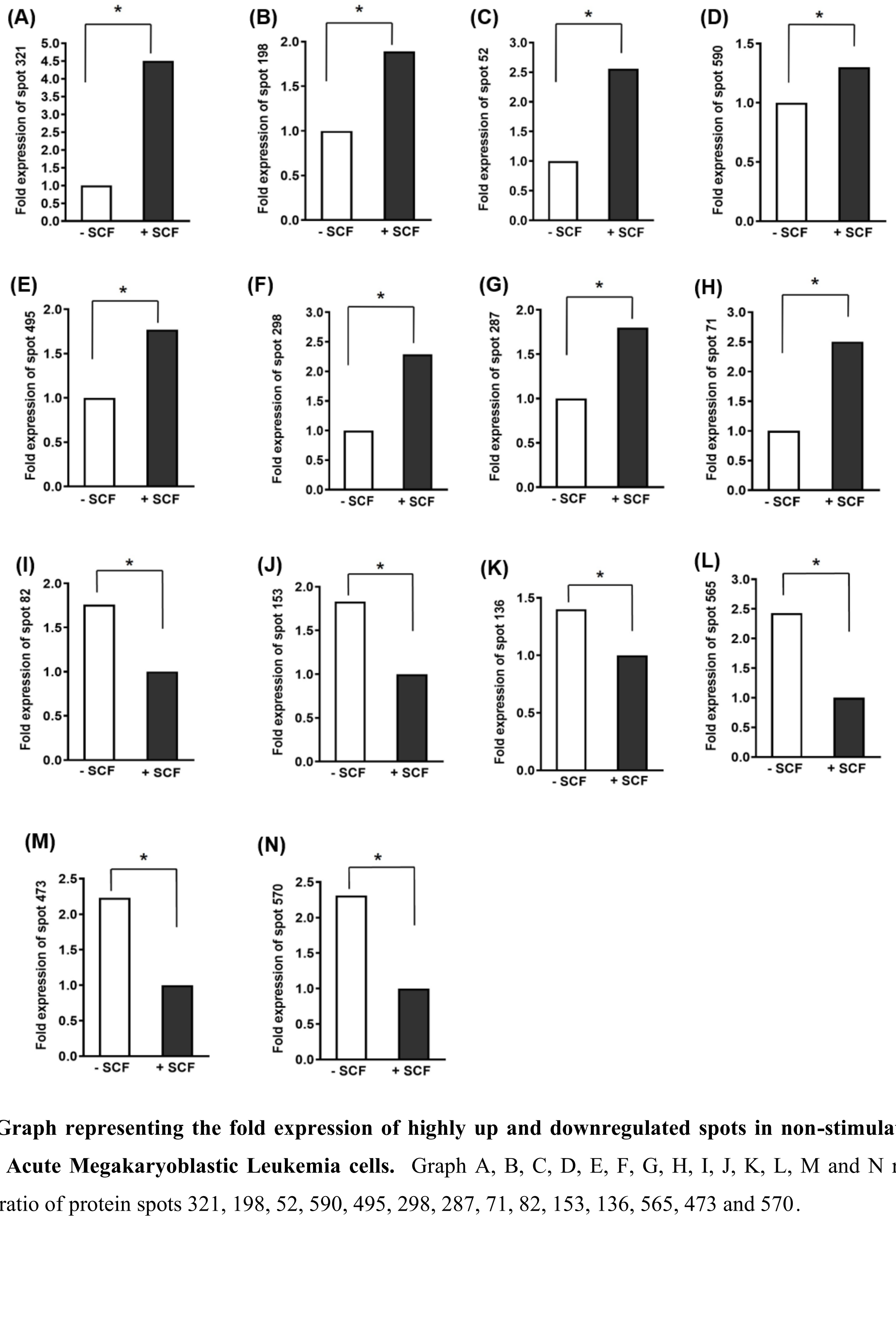


**Fig. (S4). Graph representing the fold expression of highly up and downregulated spots in non-stimulated and SCF stimulated Acute Megakaryoblastic Leukemia cells.** Graph A, B, C, D, E, F, G, H, I, J, K, L, M and N represent fold expression ratio of protein spots 321, 198, 52, 590, 495, 298, 287, 71, 82, 153, 136, 565, 473 and 570.

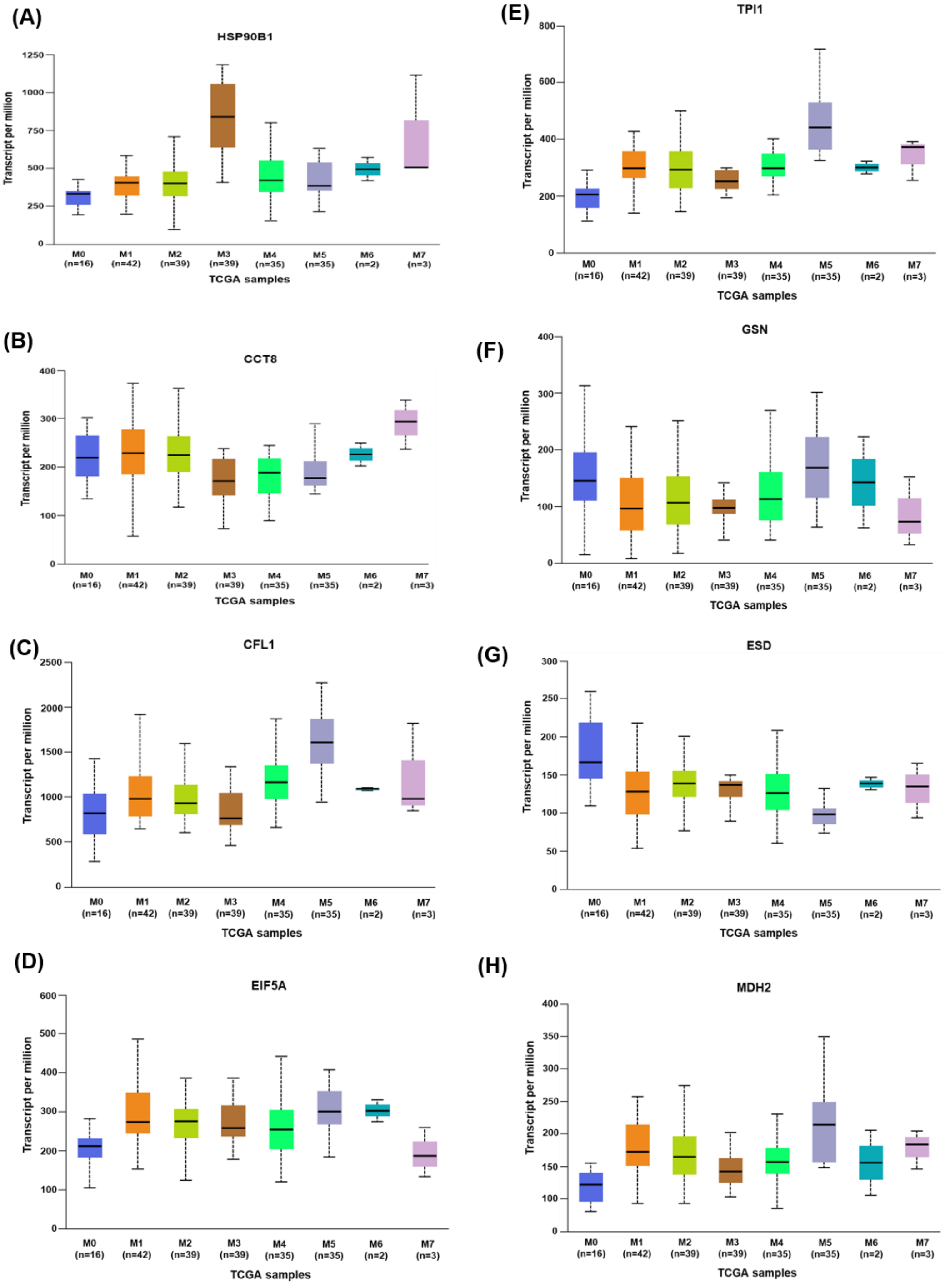


**Fig. (S5). Gene expression analysis of SCF-activated proteins across subtypes of acute myeloid leukemia (AML).** Gene expression profiles of KIT-activated proteins were analyzed in human megakaryocytic leukemia cells across different AML subtypes using UALCAN software. The analysis illustrates the differential expression patterns of selected genes among AML subgroups relative to normal controls. Data are presented as normalized expression values, highlighting subtype-specific variations. Graph A, B, C, D, E, F, G, and H represent fold expression of genes HSP90B1, CCT8, CFL1, EIF5A, TPI1, GSN, ESD and MDH2.

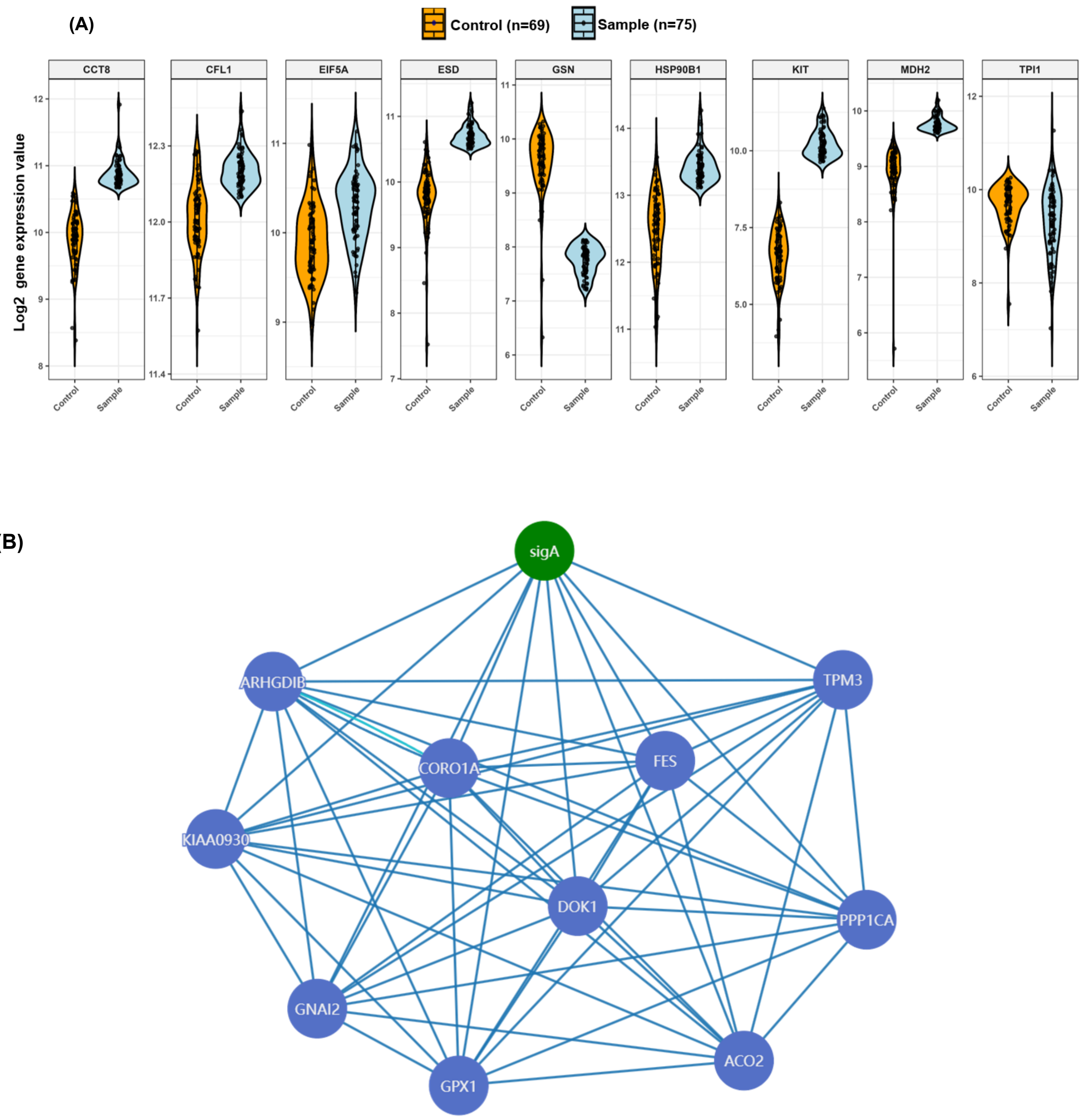


**Fig. (S6). Gene expression analysis of SCF-activated proteins in an external AML dataset.** (A) Gene expression data from the external validation dataset GSE15061, comprising 202 AML patients and 69 healthy donors, were obtained from the Gene Expression Omnibus (GEO) database. Following data normalization, the expression patterns of genes encoding the differentially expressed SCF-activated proteins were visualized using violin plots generated with the ggplot2 package in R. (B) Co-expression network of gene signature (CFL1, GSN and CCT8) with known AML prognostic genes.

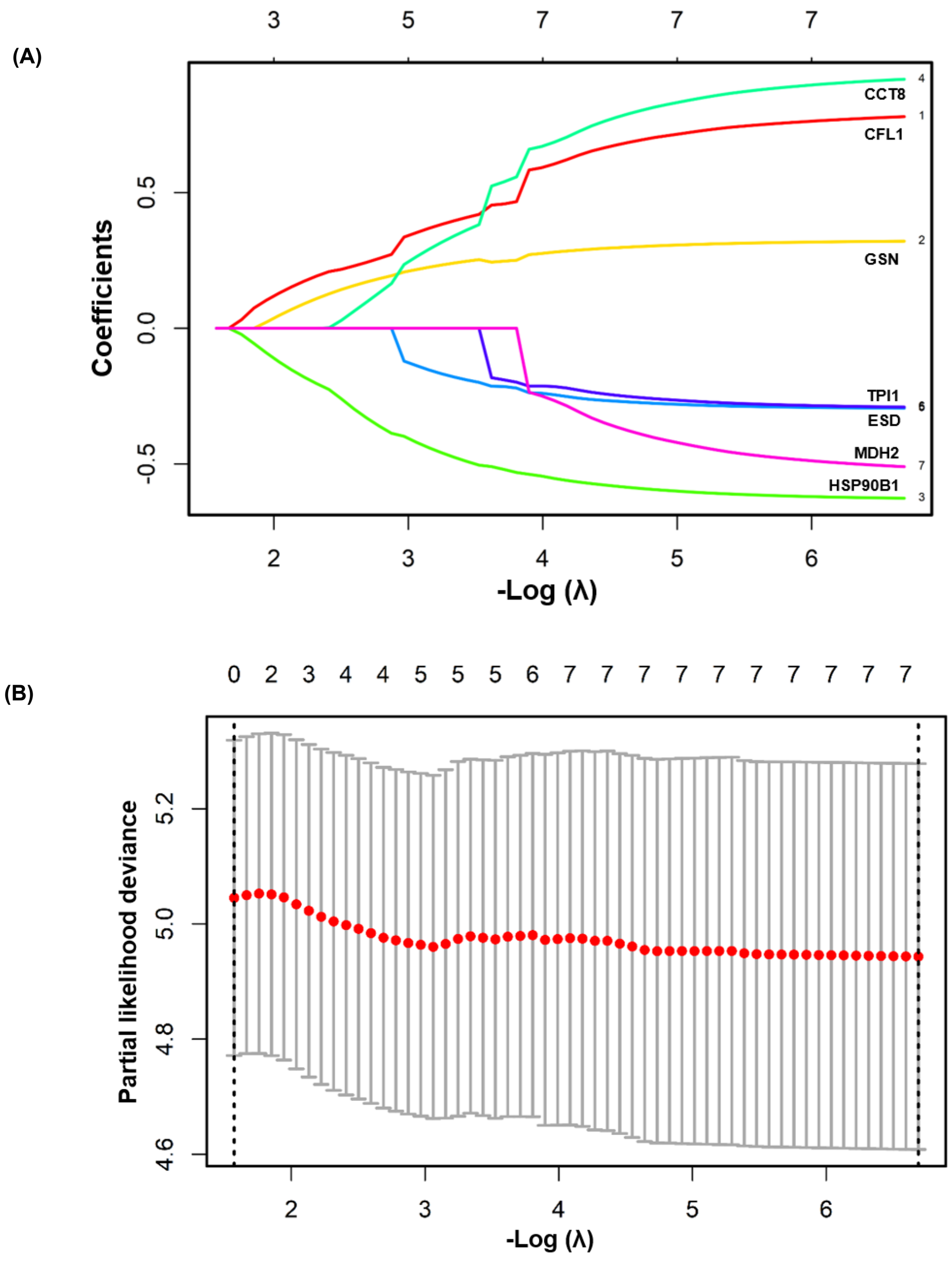


**Fig. (S7). LASSO coefficient profiles of SCF-activated LAML genes in Acute Megakaryoblastic Leukemia cells**. (A) LASSO regression analysis was performed to evaluate the contribution of genes (CFL1, GSN, CCT8, TPI, HSP90B1, ESD and MDH2) to the predicted cox-hazard model. The plot shows the coefficient profiles of each gene as a function of the regularization parameter (−log(λ)). Each colored line represents an individual gene, and the y-axis indicates the corresponding regression coefficient. As the penalty parameter increases, coefficients of less contributory genes shrink toward zero, whereas genes with stronger predictive power retain higher coefficient values. (B) Ten-fold cross-validation plot showing partial likelihood deviance as a function of log(λ). The dotted vertical lines indicate the optimal λ values, including λ.min (minimum deviance) and λ.1se (one standard error from the minimum). The optimal λ was selected to balance model performance and complexity.

**(A)**

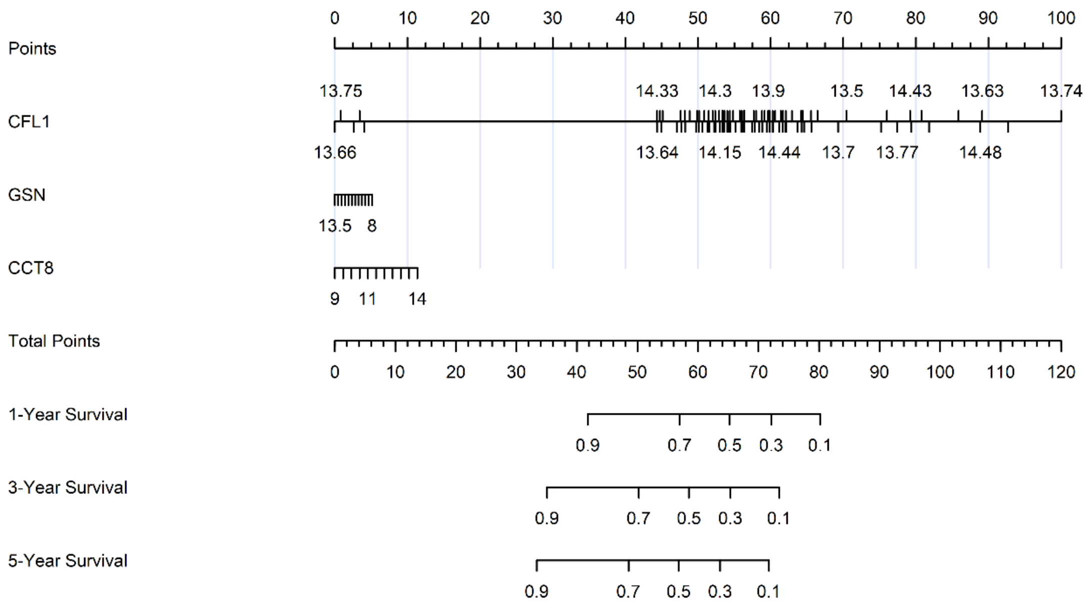


**(B)**

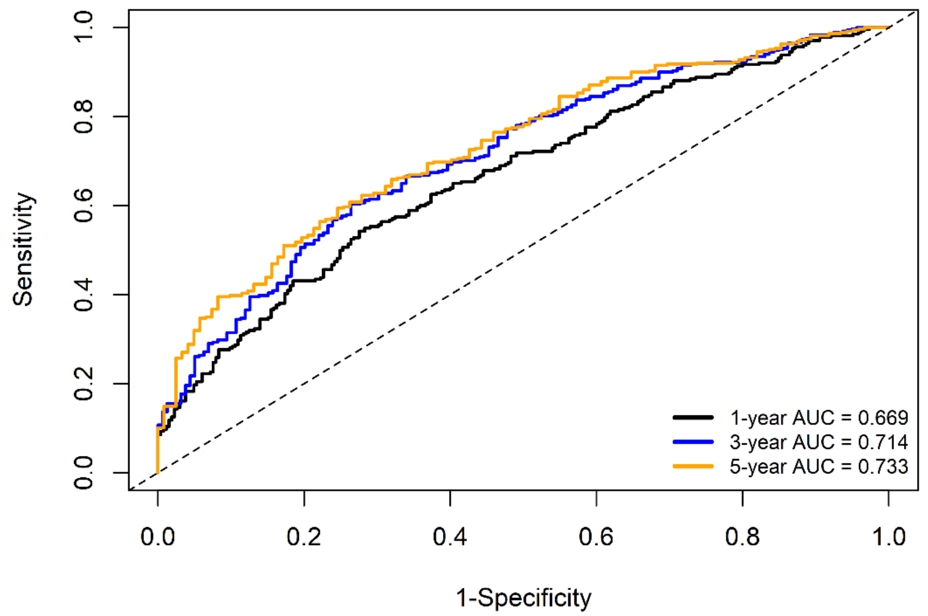


**Fig. (S8). Validation of prognostic survival model using external AML datasets**. (A) Prognostic nomogram based on AML data set GSE37642, (B) Time-ROC analysis of prognostic nomogram model.

(A)

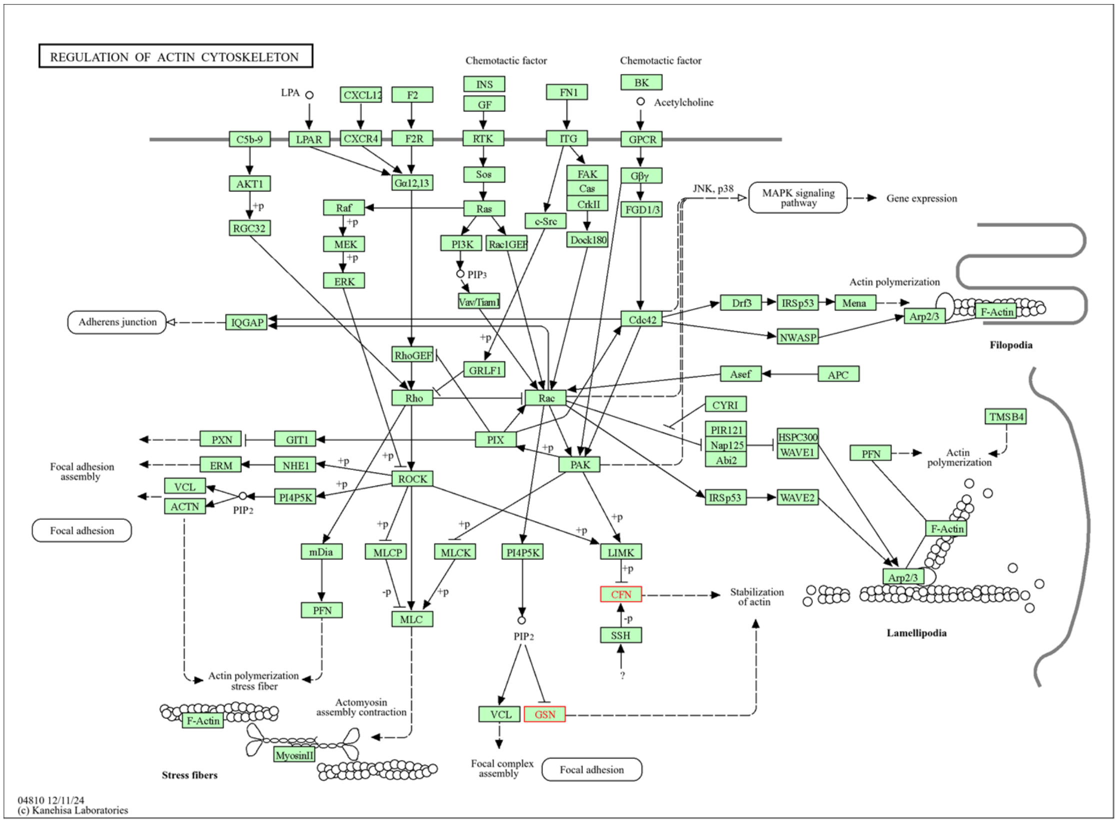
REGULATION OF ACTIN CYTOSKELETON
Chemotactic factor
Chemotactic factor
LPA
CXCL12
F2
INS
GF
FN1
BK
Acetylcholine
C5b-9
LPAR
CXCR4
F2R
RTK
ITG
GPCR
AKT1
Gα12,13
Sos
FAK
Cas
CrkII
Gβγ
JNK, p38
MAPK signaling pathway
Gene expression
RGC32
Raf
Ras
c-Src
FGD1/3
MEK
PI3K
Rac1GEF
Dock180
ERK
PIP3
Vav/Tiam1
Actin polymerization
Drf3
IRSp53
Mena
Arp2/3
F-Actin
Filopodia
Adherens junction
IQGAP
Cdc42
NWASP
RhoGEF
GRLF1
Asef
APC
Rho
Rac
CYRI
PIR121
Nap125
Abi2
HSPC300
WAVE1
TMSB4
PXN
GIT1
PIX
PAK
PFN
Actin polymerization
Focal adhesion assembly
ERM
NHE1
ROCK
IRSp53
WAVE2
VCL
ACTN
PIP2
PI4P5K
F-Actin
Focal adhesion
mDia
MLCP
MLCK
PI4P5K
LIMK
Arp2/3
CFN
Stabilization of actin
PFN
MLC
SSH
Lamellipodia
PIP2
?
Actin polymerization stress fiber
Actomyosin assembly contraction
VCL
GSN
F-Actin
MyosinII
Stress fibers
Focal complex assembly
Focal adhesion
04810 12/11/24
(c) Kanehisa Laboratories

(B)

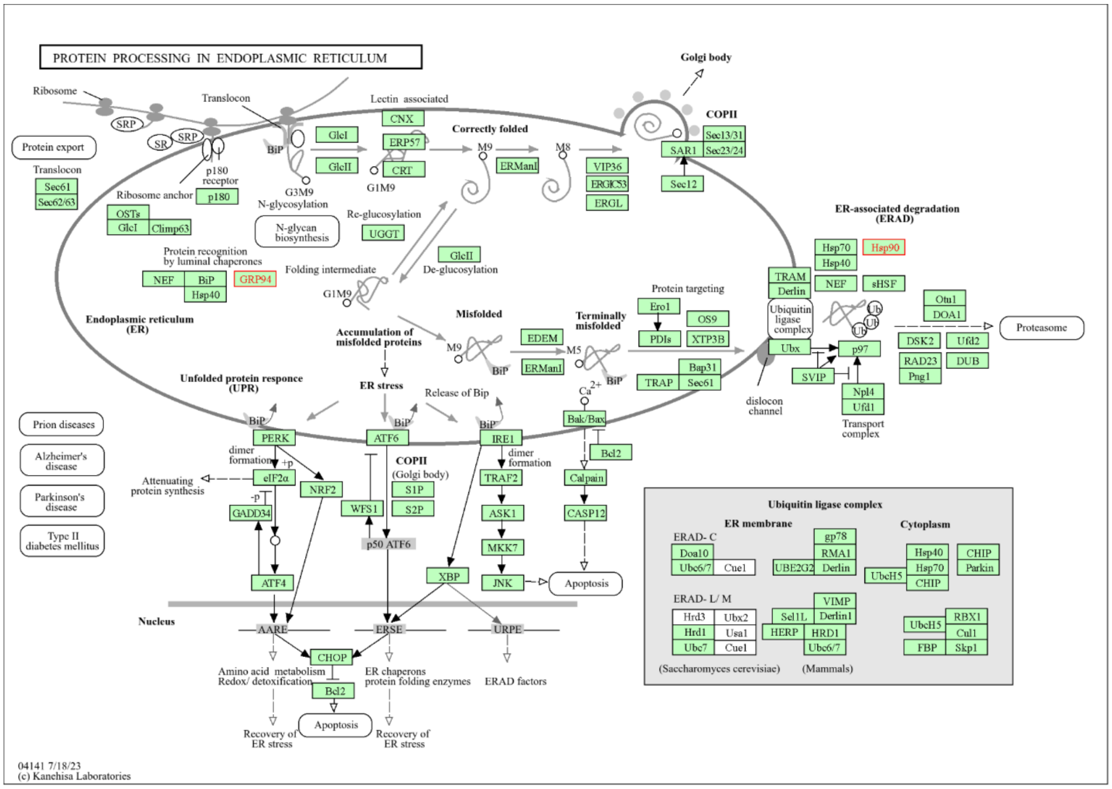
PROTEIN PROCESSING IN ENDOPLASMIC RETICULUM
Golgi body
Ribosome
Translocon
Lectin associated
COPII
SRP
SR
SRP
CNX
Correctly folded
Sec13/31
Protein export
BiP
GlcI
ERP57
M9
M8
SAR1
Sec23/24
Translocon
p180 receptor
GlcII
CRT
ERManI
VIP36
Sec61
Sec62/63
G3M9
G1M9
ERGIC53
Sec12
N-glycosylation
ERGL
Ribosome anchor
p180
Re-glucosylation
OSTs
GlcI
Climp63
N-glycan biosynthesis
UGGT
ER-associated degradation (ERAD)
Protein recognition by luminal chaperones
GlcII
De-glucosylation
Hsp70
Hsp90
Hsp40
Folding intermediate
NEF
BiP
GRP94
Hsp40
TRAM
Derlin
NEF
sHSF
Protein targeting
G1M9
Ero1
Otu1
DOA1
Ubiquitin ligase complex
Ub
Endoplasmic reticulum (ER)
Misfolded
Terminally misfolded
OS9
PDIs
XTP3B
Proteasome
Accumulation of misfolded proteins
EDEM
M9
M5
Ubx
p97
DSK2
Ufd2
RAD23
DUB
Png1
BiP
ERManI
BiP
Bap31
TRAP
Sec61
SVIP
Unfolded protein responce (UPR)
ER stress
Ca2+
Release of Bip
Npl4
Ufd1
dislocon channel
Transport complex
Prion diseases
Bak/Bax
BiP
BiP
BiP
PERK
ATF6
IRE1
Bcl2
Alzheimer's disease
dimer formation
dimer formation
Attenuating protein synthesis
eIF2α
NRF2
COPII (Golgi body)
TRAF2
Calpain
Parkinson's disease
S1P
GADD34
WFS1
S2P
ASK1
CASP12
Ubiquitin ligase complex
ER membrane
Cytoplasm
Type II diabetes mellitus
ERAD- C
gp78
Doa10
RMA1
Hsp40
CHIP
Ubc6/7
Cue1
UBE2G2
Derlin
Hsp70
Parkin
UbcH5
CHIP
p50 ATF6
MKK7
ATF4
XBP
JNK
Apoptosis
ERAD- L/ M
VIMP
Hrd3
Ubx2
Sel1L
Derlin1
RBX1
UbcH5
Nucleus
Hrd1
Usa1
HERP
HRD1
Cul1
Ubc7
Cue1
Ubc6/7
FBP
Skp1
AARE
ERSE
URPE
CHOP
(Saccharomyces cerevisiae)
(Mammals)
Amino acid metabolism
Redox/ detoxification
ER chaperons protein folding enzymes
ERAD factors
Bcl2
Apoptosis
Recovery of ER stress
Recovery of ER stress
04141 7/18/23
(c) Kanehisa Laboratories

**(C)**

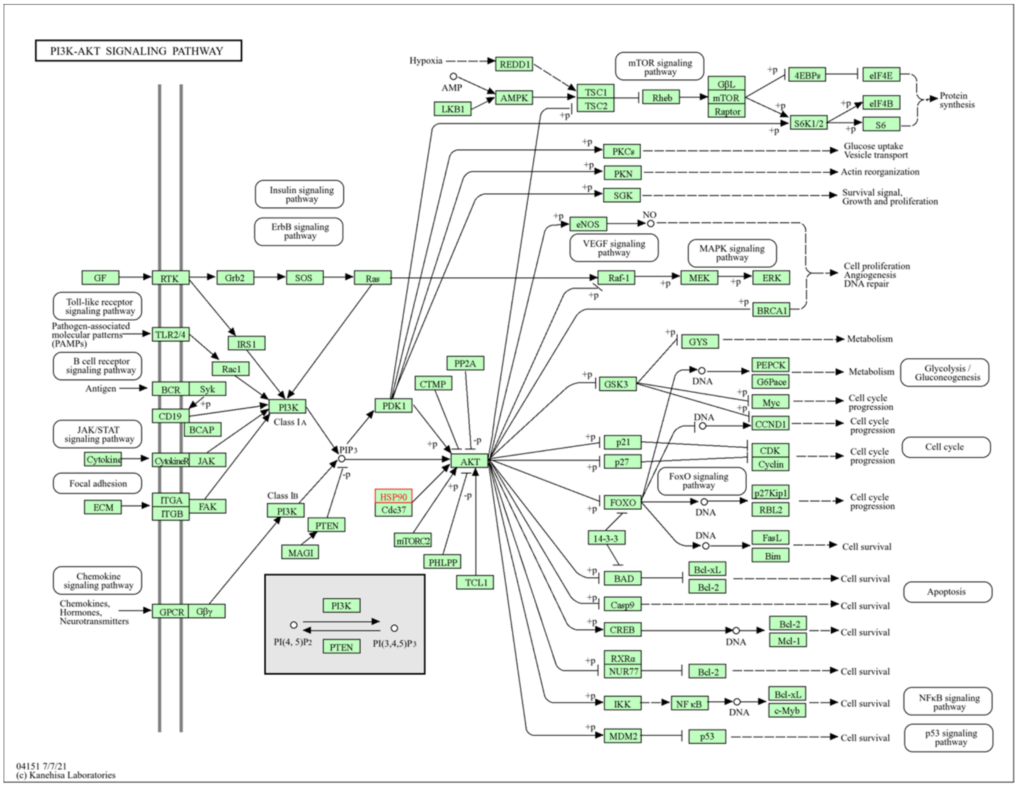

PI3K-AKT SIGNALING PATHWAY
Hypoxia
REDD1
AMP
LKB1
AMPK
TSC1
TSC2
mTOR signaling pathway
Rheb
GβL
mTOR
Raptor
4EBPs
eIF4E
eIF4B
S6K1/2
S6
Protein synthesis
PKCs
Glucose uptake
Vesicle transport
PKN
Actin reorganization
SGK
Survival signal, Growth and proliferation
Insulin signaling pathway
ErbB signaling pathway
eNOS
NO
VEGF signaling pathway
MAPK signaling pathway
Cell proliferation
Angiogenesis
DNA repair
GF
RTK
Grb2
SOS
Ras
Raf-1
MEK
ERK
BRCA1
Toll-like receptor signaling pathway
Pathogen-associated molecular patterns (PAMPs)
TLR2/4
IRS1
Rac1
B cell receptor signaling pathway
Antigen
BCR
Syk
CD19
BCAP
PI3K
Class IA
JAK/STAT signaling pathway
Cytokine
CytokineR
JAK
Focal adhesion
ECM
ITGA
ITGB
FAK
PIP3
Class IB
PTEN
MAGI
PP2A
CTMP
PDK1
AKT
HSP90
Cdc37
mTORC2
PHLPP
TCL1
GYS
Metabolism
GSK3
DNA
PEPCK
G6Pace
Glycolysis / Gluconeogenesis
Myc
CCND1
Cell cycle progression
p21
p27
CDK
Cyclin
Cell cycle
FoxO signaling pathway
FOXO
p27Kip1
RBL2
14-3-3
FasL
Bim
Cell survival
BAD
Bcl-xL
Bcl-2
Casp9
Apoptosis
CREB
Mcl-1
RXRα
NUR77
IKK
NF κB
c-Myb
NFκB signaling pathway
MDM2
p53
p53 signaling pathway
Chemokine signaling pathway
Chemokines, Hormones, Neurotransmitters
GPCR
Gβγ
PI(4, 5)P2
PI(3,4,5)P3
04151 7/7/21
(c) Kanehisa Laboratories


**(D)**

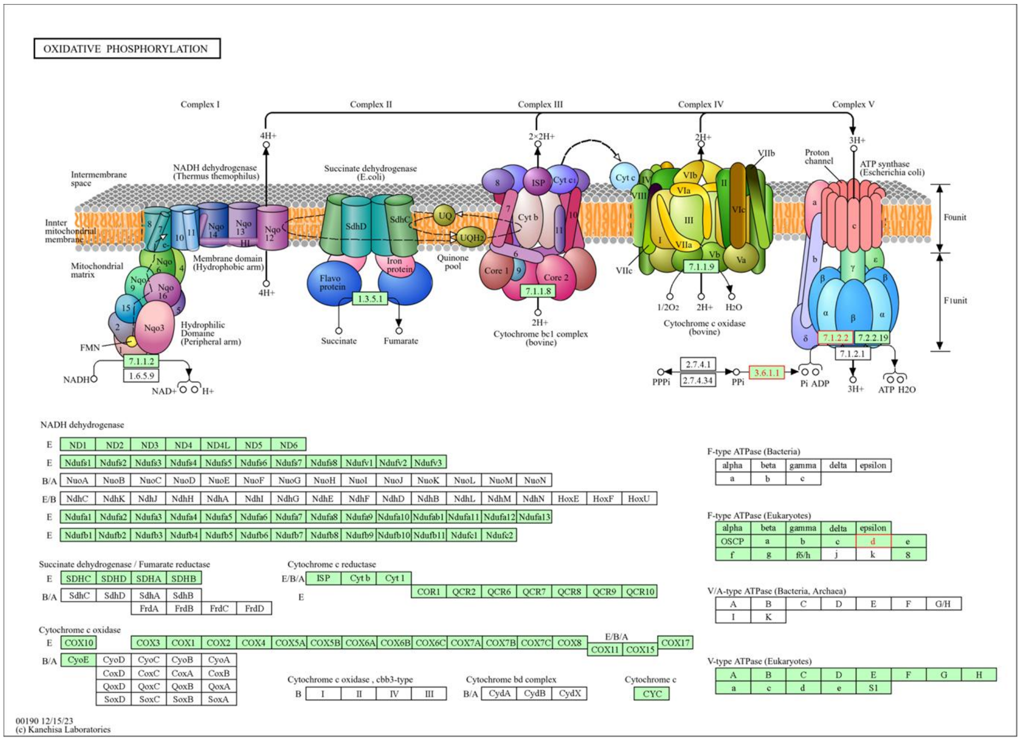

OXIDATIVE PHOSPHORYLATION
Complex I
Complex II
Complex III
Complex IV
Complex V
4H+
2×2H+
2H+
3H+
Intermembrane space
Innter mitochondrial membrane
Mitochondrial matrix
NADH dehydrogenase (Thermus themophilus)
Membrane domain (Hydrophobic arm)
Hydrophilic Domaine (Peripheral arm)
FMN
NADH
NAD+
H+
7.1.1.2
1.6.5.9
Succinate dehydrogenase (E.coli)
SdhD
SdhC
Flavo protein
Iron protein
1.3.5.1
Succinate
Fumarate
UQ
UQH2
Quinone pool
ISP
Cyt c1
Cyt b
Core 1
Core 2
7.1.1.8
Cytochrome bc1 complex (bovine)
Cyt c
VIb
VIa
VIIa
VIIc
Vb
Va
VIc
7.1.1.9
1/2O2
H2O
Cytochrome c oxidase (bovine)
Proton channel
ATP synthase (Escherichia coli)
Founit
F1unit
7.1.2.2
7.2.2.19
7.1.2.1
2.7.4.1
2.7.4.34
3.6.1.1
PPPi
PPi
Pi ADP
ATP H2O
NADH dehydrogenase
E ND1 ND2 ND3 ND4 ND4L ND5 ND6
E Ndufs1 Ndufs2 Ndufs3 Ndufs4 Ndufs5 Ndufs6 Ndufs7 Ndufs8 Ndufv1 Ndufv2 Ndufv3
B/A NuoA NuoB NuoC NuoD NuoE NuoF NuoG NuoH NuoI NuoJ NuoK NuoL NuoM NuoN
E/B NdhC NdhK NdhJ NdhH NdhA NdhI NdhG NdhE NdhF NdhD NdhB NdhL NdhM NdhN HoxE HoxF HoxU
E Ndufa1 Ndufa2 Ndufa3 Ndufa4 Ndufa5 Ndufa6 Ndufa7 Ndufa8 Ndufa9 Ndufa10 Ndufab1 Ndufa11 Ndufa12 Ndufa13
E Ndufb1 Ndufb2 Ndufb3 Ndufb4 Ndufb5 Ndufb6 Ndufb7 Ndufb8 Ndufb9 Ndufb10 Ndufb11 Ndufc1 Ndufc2
Succinate dehydrogenase / Fumarate reductase
E SDHC SDHD SDHA SDHB
B/A SdhC SdhD SdhA SdhB
FrdA FrdB FrdC FrdD
Cytochrome c reductase
E/B/A ISP Cyt b Cyt 1
E COR1 QCR2 QCR6 QCR7 QCR8 QCR9 QCR10
Cytochrome c oxidase
E COX10 COX3 COX1 COX2 COX4 COX5A COX5B COX6A COX6B COX6C COX7A COX7B COX7C COX8 COX11 COX15 COX17
B/A CyoE CyoD CyoC CyoB CyoA
CtaB CoxD CoxC CoxA CoxB
QoxD QoxC QoxB QoxA
SoxD SoxC SoxB SoxA
Cytochrome c oxidase , cbb3-type
B I II IV III
Cytochrome bd complex
B/A CydA CydB CydX
Cytochrome c
CYC
F-type ATPase (Bacteria)
alpha beta gamma delta epsilon
a b c
F-type ATPase (Eukaryotes)
alpha beta gamma delta epsilon
OSCP a b c d e
f g f6/h j k 8
V/A-type ATPase (Bacteria, Archaea)
A B C D E F G/H
I K
V-type ATPase (Eukaryotes)
A B C D E F G H
a c d e S1
00190 12/15/23
(c) Kanehisa Laboratories

**Fig. (S9). KEGG Signalling pathway analysis of differentially expressed protein in SCF stimulated acute Megakaryoblastic Leukemia cells.** (A) Regulation of Actin Cytoskeleton, (B) Protein processing in Endoplasmic Reticulum, (C) PI3K-AKT Signalling, (E) Oxidative phosphorylation and (D) Lipid and Atherosclerosis.
